# KG-Microbe - Building Modular and Scalable Knowledge Graphs for Microbiome and Microbial Sciences

**DOI:** 10.1101/2025.02.24.639989

**Authors:** Brook E. Santangelo, Harshad Hegde, J. Harry Caufield, Justin Reese, Tomas Kliegr, Lawrence E. Hunter, Catherine A. Lozupone, Christopher J. Mungall, Marcin P. Joachimiak

## Abstract

The integration of many disparate forms of data is essential for understanding the microbial world and its interaction with the environment and human health. Doing so is particularly challenging in the context of microbe-host and microbe-microbe interactions that contribute to health or environmental outcomes. There are often thousands of relevant microbial species, and millions of interactions among those microbes and with their environment or host. Some experimental observations only distinguish coarser taxonomic resolutions such as family or phylum-level. Integrated information (e.g., about host and microbial physiology, genetics, and metabolism) facilitates deeper understanding of complex interactions and helps interpret correlative results. The KG-Microbe construction framework is a novel approach to harmonizing bacterial and archaeal data in the form of a knowledge graph (KG). Starting from a core KG with organismal traits, environments and growth preferences, the framework generates a hierarchy of related KGs targeting specific conceptual use cases, including the human host-associated microbiome in the context of disease. KG-Microbe is a standardized and interoperable framework that integrates microbial organismal and genomic traits, represented ontologically, for biomedical, environmental, and other applications. The framework supports customizable taxa subsets representing microbial lineages or communities of interest. Evaluations of the KG-Microbe knowledge graphs through a series of competency questions demonstrate the accuracy and effectiveness of the data harmonization, and the utility of the resulting KGs in inflammatory bowel and Parkinson’s diseases. Finally, the predictive and environmental capabilities of the KGs are demonstrated by explaining growth preferences through training a model using graph features. KG-Microbe is a flexible, modular enabling technology for humans and machine learning methods to uncover mechanistic explanations of microbial associations.

## 1. Introduction

Our knowledge of the influence of microbes in varied environments has expanded significantly due to technological advances. The ability to identify taxa using efficient sequencing technologies has allowed us to survey many environments and hosts with respect to differences in populations of microbial taxa and their associated molecular functions, such as understanding the gut microbiome of a disease versus a healthy population^1,2^ and the biology of the constituent microbes

Sequence-based methods for understanding microbial function either only cover a specific set of taxa or predict function at potentially coarse taxonomic resolution, such as the family or phylum level^1,3^. Attributing functions at more specific taxonomic resolution can be done by considering taxonomic relationships (e.g., between species and strains) between annotated and unannotated microbes. For example, one critical function of microbes with implications for health and disease is the production of short-chain fatty acids (SCFAs). Evidence that these microbial metabolites reduce inflammation suggests the protective nature of SCFA-producing microbes against inflammatory disease^4^. Understanding this function at a per-strain or species level is important for characterizing a healthy microbiome and distinguishing it from disease states, yet difficult given the fragmented nature of microbial knowledge.

Information about the structure and function of microbes and their communities is available in a range of publicly available sources. Microbial traits and functional attributes can be discovered with laboratory experiments or computational predictions using publicly available data. However, relevant information is spread across many special-purpose databases, which vary greatly in structure and format. A knowledge graph (KG) is an effective and flexible way to integrate this vast amount of information, consistently representing knowledge despite varying formats, sources, and domains of the original data sources. KGs often rely on ontologies, which are expert curated, hierarchical summaries of concepts in a specific domain, e.g. biological metabolites, that support standardized knowledge representation. In the biomedical sciences, KGs have gained popularity in applications such as drug discovery and repurposing and disease classification^5,6^. The inherent data interoperability of KGs presents a unique benefit to the study of microbial communities as a rapidly evolving field that is still undergoing harmonization. Applying a KG approach to microbiome related research questions is in its early stages, as existing microbial KGs represent targeted aspects of microbiology or human disease^7–10^. These resources have demonstrated the potential for KGs to enhance microbiome research through answers to through queries for broad biomedically and environmentally relevant information. However, no resource exists that harmonizes microbial traits and genomic annotations while seamlessly integrating taxonomic relationships.

To address this gap, we developed the KG-Microbe framework which supports the generation of large and comprehensive KGs with relevant information about microbes and their interactions with the environment and human health. The KG-Microbe framework can ingest data from a broad array of sources, ranging from ontologies to published datasets to reference databases, to fully represent these interactions. KG-Microbe promises to enhance microbial and microbiome research through representation of microbial morphology, traits, ecology, function, and taxonomy. As very large KGs can be computationally demanding to exploit, the KG-Microbe framework enables the construction of a set of KGs, fit for purpose in varied contexts by including only relevant information for a particular use case. The quality of KG-Microbe KGs is demonstrated through graph competency queries, which showcase the variety of data sources and the relevance of content within the graph, and its effectiveness is demonstrated through its ability to enhance our functional understanding of associations with inflammatory bowel disease (IBD) and Parkinson’s disease (PD). We also demonstrate the utility of KG-Microbe in environmental contexts, using KG features for training machine learning models to predict and explain temperature growth preferences, leading to new biological insights. The KG-Microbe framework can enable the microbiology and microbiome fields by harmonizing and standardizing microbial data, facilitating new predictions about microbes and their communities.

## 2. Results

### 2.1. KG-Microbe Content

The KG-Microbe framework supports the integration of microbial trait and function data from curated databases, ontologies, and experimental results. Because computation and memory are limiting factors, the framework supports the construction of custom KGs that synthesize data sources relevant to applications of microbiology or biomedicine, otherwise known as modules (Figure 1). The core KG, KG-Microbe-Core, designed to contain a foundational set of broadly applicable knowledge, includes microbial organismal traits obtained from structured databases, which sourced data from literature curation or experimental assay results. KG-Microbe-Biomedical introduces disease relevant concepts and human functional information. KG-Microbe-Function and KG-Microbe-Biomedical-Function also ingest microbial functional information (Figure 1A). All KG-Microbe graphs are structured according to the Biolink Model, a formalized data model for the representation of complex biological data^11^. The KG-Microbe framework provides software infrastructure to build the four main graphs, which are made available in the KG-Registry^12^. Knowledge of microbial traits and functions was ingested from a variety of resources and modeled to serve the purposes of mechanistic inference in different research domains (Figure 2).

**Figure 1.**
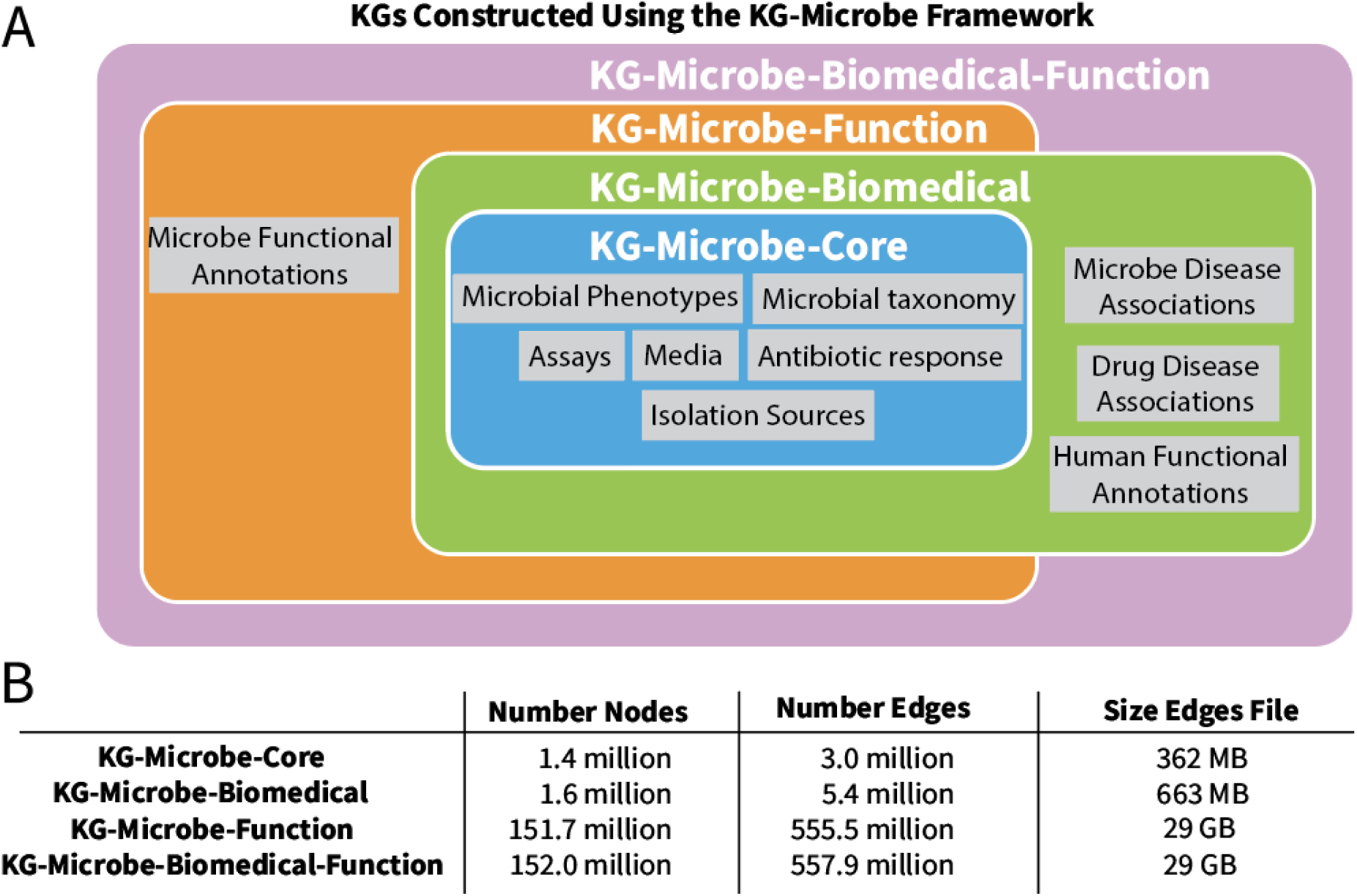
(A) Content of each KG generated with the KG-Microbe framework. (B) The size of each KG, listed as total number of nodes, total number of edges, and the size of the edges file. Edge file sizes are listed according to the SUSE Linux Enterprise Server 15 SP5 system. Each KG in the framework can be used as a starting point to construct novel custom KGs.

**Figure 2.**
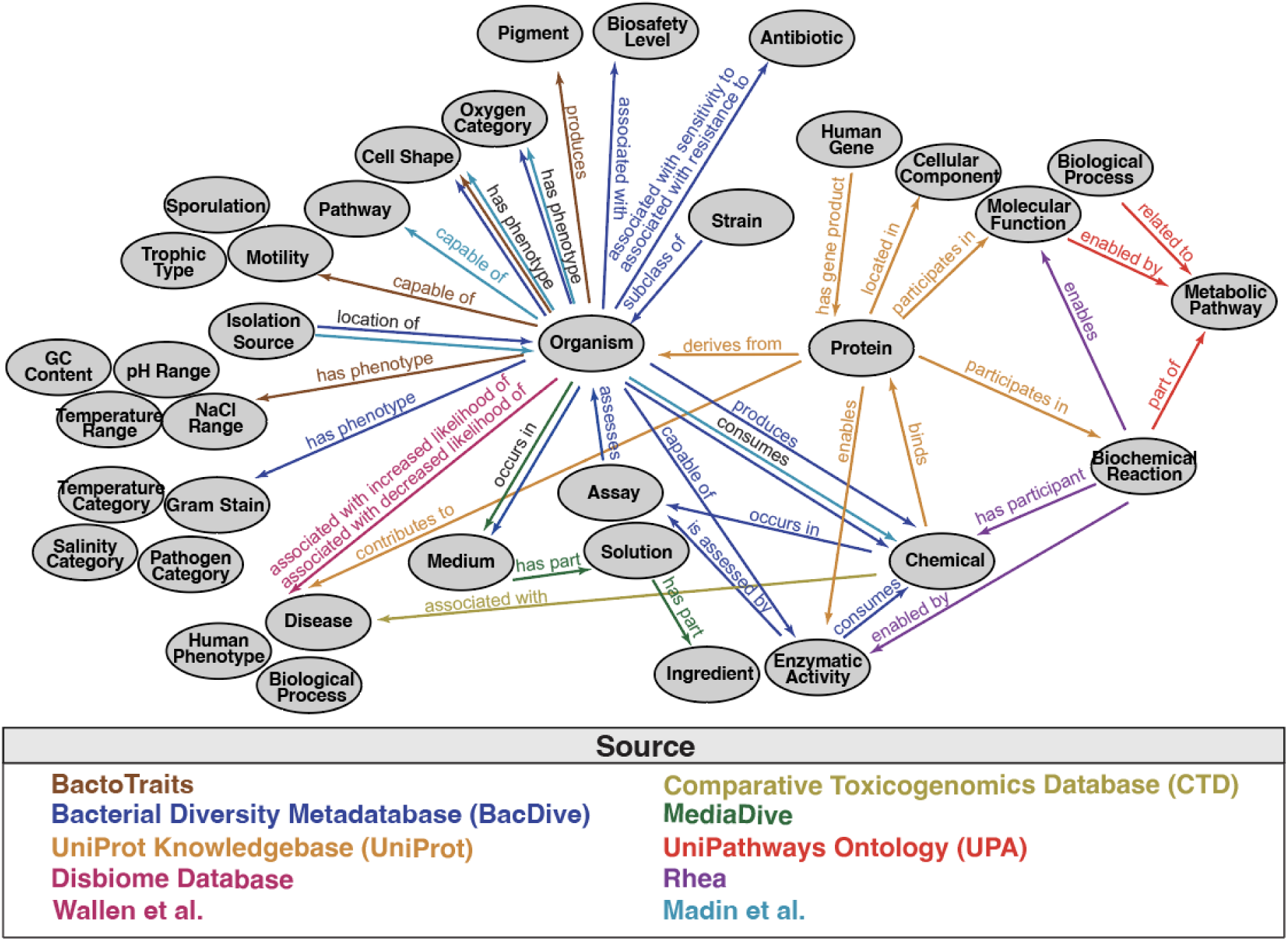
Semantic schema supported by the KG-Microbe framework, representing all sources across all KGs in the KG-Microbe framework.

#### 2.1.1. Microbial Taxonomy and Strain Representation

The value of KGs produced by the KG-Microbe framework depends on the accurate representation of microbial knowledge at all taxonomic levels. KG-Microbe-Core includes 1,012,214 microbial taxa nodes, 870,981 of which are normalized to National Center for Biotechnology Information (NCBI) Taxonomy terms^13^. The remaining 159,680 taxa without a corresponding NCBI Taxonomy term are introduced as children of their related species level NCBI Taxonomy term with the *“strain”* prefix (using the predicate *“biolink:subclass_of”*), for example *Faecalibacterium prausnitzii L2-6*, a strain, is a subclass of *Faecalibacterium prausnitzii,* a species. The inclusion of the full taxonomic hierarchy supports summarization of traits at higher taxonomic levels or exploration of traits at lower taxonomic levels, using the structured relationships in the taxonomic hierarchy.

Traits and genomic annotations are linked to microbial nodes at a species or strain level. Strains are normalized to NCBI Taxonomy when a mapping to an NCBI Taxonomy node exists in the source. This sometimes results in a rollup of these attributes to the species level, where multiple strain level traits are attached to a species level node. In these cases, the strain nodes are still introduced to the KG as children of their related species, and the provenance of that trait-strain relationship remains in the graph metadata, for example *“strain:CCUG-59342”*, a strain, is a subclass of *Escherichia coli (NCBITaxon:562)*, the species to which all traits are linked. When no mapping to an NCBI Taxonomy node exists in the source, traits are linked directly to a corresponding strain level node in the KG. This modeling choice supports the consistency and robustness of taxa representation, with traits linked to standardized terms for microbial data that is not strain-resolved by identifier. This also supports efficient queries when mapping external data sources to the knowledge represented in the graph.

#### 2.1.2. Microbial Phenotypes

Microbial phenotypes are ingested from three sources: the Bacterial Diversity Metadatabase (BacDive), the BactoTraits database, or a published dataset that unifies various sources of information about microbiology Madin et al.^14–16^ The Bacterial Diversity Metadatabase (BacDive) is one of the largest standardized resources of prokaryotic information from over 80,000 strains^17^. The BactoTraits database expands on BacDive cultured bacteria experimental results by modeling 19 physiologic and morphological traits^15^. Madin et al. is a merged dataset of over 170,000 strains and 15,000 species curated from a wide range of published sources including publications and textbooks^16^. Each of the microbial phenotype nodes are attached to a species or strain level taxon node (Figure 2).

Environmental growth information including four categories of temperature preference (e.g. *“mesophilic”*), 9 categories of oxygen preference (e.g. *“microaerophile”*), and 7 categories of salinity (e.g. *“halophilic”*), each of which are introduced as unique categorical nodes with prefixes that standardize the node type (see Methods). Specific growth or survival conditions based on quantitative results of temperature, pH, and NaCl concentration are also introduced including the optimal range, the growth-permissible range, and ecological spread for growth.

Morphological qualities of microbes introduced to the KG include four bins of the percentage of GC content in the genome (e.g. *“high”* or *“low”*), 32 cell shapes (e.g. *“rod”*), four bins of cell length and width (e.g. *“high”* or *“low”*), and whether a microbe is spore-forming or motile.

Several forms of microbial metabolism are modeled in the KG. Direct microbe-chemical links indicate whether a microbe is a producer or consumer of a given chemical. When ingested from BacDive or Madin et al., this knowledge comes from culture observations of carbon substrates or published literature, serving as a developing gold-standard of microbial metabolism. 29 categories of trophic type (e.g., *“autotrophy*) are also included. Antibiotic resistance and sensitivity data is sourced from antibiogram results in BacDive. All antibiotic compounds are mapped to ChEBI nodes, which consist of 165 unique chemicals to which microbes are sensitive to and 155 unique chemicals to which microbes are resistant (e.g. *“chloramphenicol”*, *“amoxicillin”*, and *“ampicillin”*). Finally, the involvement of species or strains in pathways or capabilities to perform enzymatic activities are introduced from culture observations or structured databases.

Ecological phenotypes such as pathogenicity in a given host (e.g. *“plant”* or *“human”*), biosafety level (BSL) and the sources from which a given strain has been found, referred to as an isolation source, are also modeled. The isolation source nodes enable the filtering of microbes from a given environment or characterization of a microbial community with links to a specific environment, for example the host large intestine.

#### 2.1.3. Medium and Assay Entities

We modeled a series of assay results called Analytical Profile Index (API) tests which assess a series of physiological traits, from Bacdive^14^. The assays from which the culturing results are sourced in Bacdive are introduced according to the microbe that each assay tests. 461 unique assays are represented with the chemical that the assay intends to test (*“chemical, biolink:occurs_in, assay”*) or the enzymatic activities assessed by the assay (*“enzymatic activity, biolink:assessed_by, assay”*). Information about media was also introduced from MediaDive, an expert curated database of over 3,000 distinct cultivation media recipes for over 40,000 strains that are linked to the strains represented by BacDive^18^. From this source, we introduce 3,333 unique media in which microbes can occur with the triple pattern “*microbe, biolink:occurs_in, medium”*. The chemical solutions that make up each medium are then modeled as unique nodes that have chemical parts (*“solution, biolink:has_part, chemical”*). All of this information is valuable to aid in understanding microbial biology in the context of nutrition requirements and growth preferences.

#### 2.1.4. Functional Annotations

Functional annotations for 32,662 microbes are ingested by the KG-Microbe framework from the UniProt Knowledgebase (UniProt).^19^ Due to its large size (Figure 1B), UniProt data is only ingested into KG-Microbe-Function and KG-Microbe-Biomedical-Function. UniProt is a leading community and expert curated database of both reviewed and computationally predicted functional annotations of proteins for a large number of organisms^19^. UniProt is updated frequently and thus provides comprehensive and up to date functional annotations critical for understanding microbial mechanisms. KG-Microbe ingests functional annotations by organism (NCBITaxon) proteomes, which is a set of proteins encoded by the genes that are present as complete or draft genome, when the proteome contains a threshold number of proteins (see Methods). The protein node is linked directly to the corresponding NCBI Taxonomy node using the pattern *“protein, biolink:derives_from, microbe”*, then linked to the chemicals to which the protein binds, biological processes or molecular functions that the protein participates in, cellular components that the protein is located in, enzymatic activity that the protein is capable of, and biochemical reactions that the protein participates in (Figure 2). Enzymatic activities are mapped to Enzymatic Commission (EC) numbers, which provide a hierarchical structure that categorizes enzymes by function. Biochemical reactions are mapped to IDs from the Rhea database, an expert-curated knowledge base of reactions and their relationships to chemical substrates and products. Molecular function and biological processes are mapped to IDs from the Gene Ontology (GO). By representing protein function across all three of these similar sources, none of the curated information from UniProt about protein function is lost. Using these standardized data resources ensures cross-database compatibility of the represented knowledge.

Pathway information is also represented using the Unipathways ontology, a manually curated resource of metabolic pathways that provides relationships between biochemical reactions, enzymatic activity, and biological processes or molecular functions^20^. In the KG, the information is abstracted to the molecular function that is enabled by the pathway, the synonym biological process for each pathway when available (linked by a *“biolink:related_to”* edge), and the biochemical reactions that are part of each pathway. This semantic representation of genome annotations, within the context of inter-linked reaction, chemical, function, and pathway hierarchies, provides a standardized way to formally define microbial protein function, where provenance is maintained by including all relevant function node types.

#### 2.1.5. Biomedically Relevant Insights

Disease and human-host relevant information was harmonized into the biomedical KG-Microbe graphs for applications in understanding the involvement of the host microbiome in disease. The human subset of functional annotations from UniProt are introduced to KG-Microbe-Biomedical and KG-Microbe-Biomedical-Function, which resulted in 163,341 new protein nodes. The gene templates of human proteins are also introduced from UniProt^21^. This supports a combined insight into host biology and microbiology in pursuit of exploring the host-associated microbiome. Human diseases and their relationships to microbes are introduced from two resources: Disbiome and a published set of differentially abundant taxa from a population study of PD, referred to as Wallen et al.^22,23^. Each source provides correlative relationships between a taxon and a disease. Disbiome is a database of published information about microbial composition in disease and health states for diseases or human phenotypes^22^. Microbes that are found to be increased in a disease gut microbiome are linked to the corresponding disease or phenotype with the predicate *“biolink:associated_with_increased likelihood_of”*, of which there are 4,209 unique associations, whereas those decreased in a disease context are linked via the *“biolink:associated_with_decreased likelihood_of”* predicate, of which there are 4,618 unique associations. Chemical-disease associations are sourced from the Comparative Toxicogenomics Database (CTD) and modeled as *“chemical, biolink:associated_with, disease”*^24^. With this human physiology and disease knowledge, KG-Microbe-Biomedical and KG-Microbe-Biomedical-Function support biomedically relevant inference.

### 2.2. Examining KG-Microbe Content through Competency Questions

The integration of vast amounts of information included in the KG-Microbe framework can include errors or omissions. We assess the accuracy of KG-Microbe-Function demonstrating four main qualities: (1) the graph’s ability to synthesize multiple sources of information, (2) the graph’s ability to infer microbial trait and function across taxonomic relationships, (3) the accessibility of complex knowledge from the graph enabled by its formal and computationally interoperable representation, and (4) the relevance of the graph content to biomedical applications of the host associated microbiome.

The intestinal mucosa is an important barrier separating commensal microbes from the host. Short chain fatty acids (SCFAs) are made from nondigestible carbohydrates by microbial fermentation in the human gut. Butyrate is one of several SCFAs that have been shown to play an important role in strengthening the intestinal barrier^25^. Butyrate participates in several main signalling pathways critical to epithelial barrier function and immune homeostasis, including the AMPK and Akt signalling pathway that facilitates tight junction assembly and the activation of G-protein-coupled receptors to induce T cell-independent IgA secretion in the colon^26^.

To understand the representation of butyrate production in KG-Microbe-Function, we evaluated specific paths, i.e. series of linked nodes and edges, in the KG that model biological mechanisms that are herein referred to as *semantic representations* (Figure 4A). By capitalizing on the unique capabilities of a KG to model diverse data types, semantic representations offer a powerful methodology for accessing knowledge. We chose three semantic representations from two sources, BacDive and UniProt, that each describe butyrate production in different ways. The first semantic representation is from BacDive organismal traits, reporting that a given microbe produces butyrate. Another semantic representation uses biochemical reactions (as Rhea nodes) that involve butyrate as a participant from UniProt protein annotations for a given microbe. To predict reaction direction and specify butyrate as a product rather than a participant, we used the eQuilibrator method (see Methods)^27^. The final semantic representation includes sets of enzymatic activity annotations (as EC nodes) for proteins from a given microbe that are involved in one of four butyrate producing pathways (the acetyl-CoA, glutarate, 4-aminobutyrate, and lysine pathways, Figure 4A), which were identical EC sets to those used in a butyrate pathway prediction analysis to which we compared the microbial taxa resulting from our competency queries (referred to as Vital et al.)^28^. While the content of the two semantic representations from UniProt is similar, we explore the differences in protein annotations with biochemical reactions (Rhea) vs. enzymatic activity (EC). The inconsistencies of the resulting butyrate-producing microbial set from each semantic representation exemplifies the need for data harmonization to capture all knowledge.

**Figure 3.**
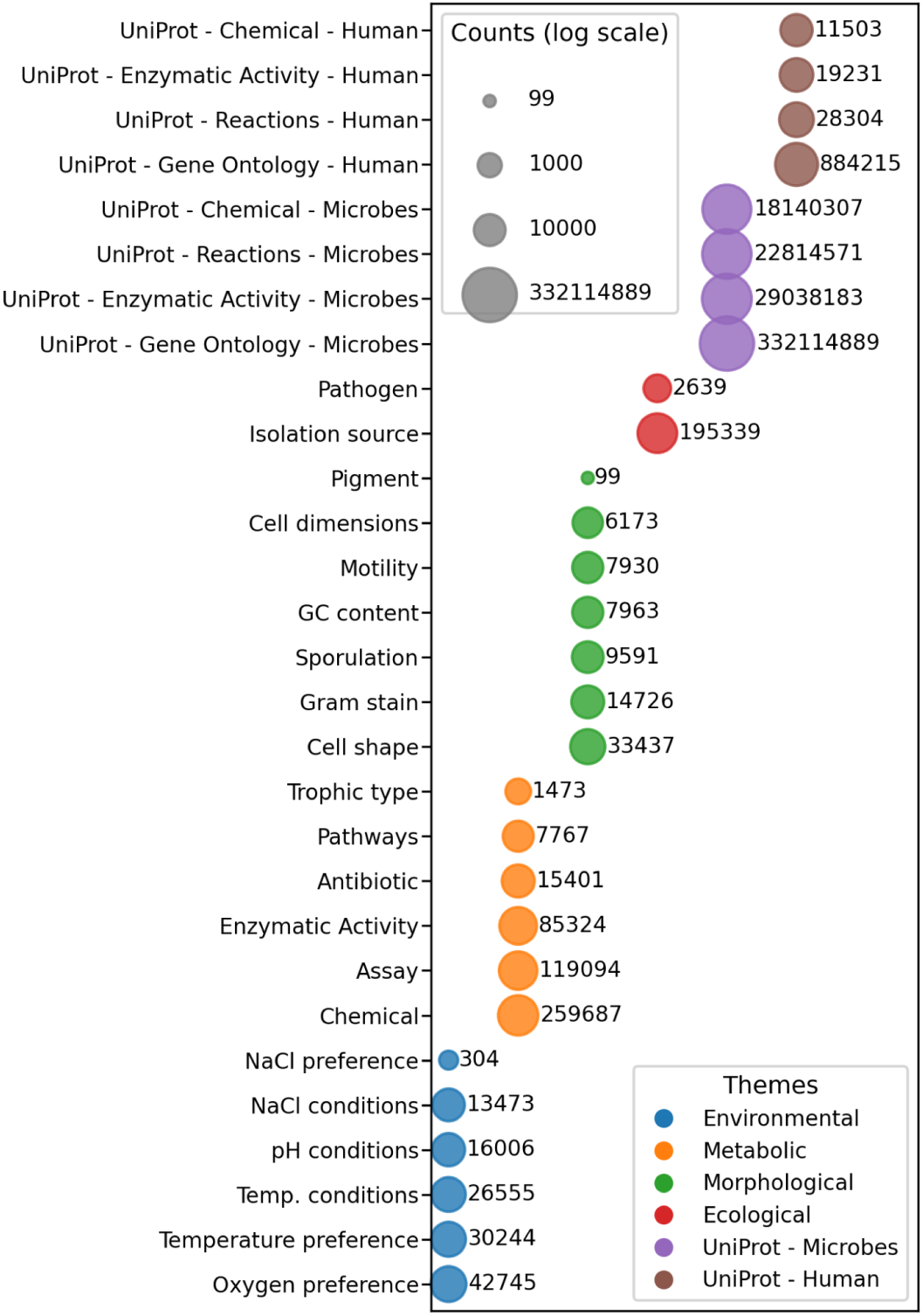
Summary of organismal traits and themes in KG-Microbe. The y-axis represents ontology or custom prefixes corresponding to different traits. Each trait prefix can have many different values and the numbers in the plot represent the count of taxa which have any value with that prefix. Traits with fewer than five edges are not shown. KG-Microbe has annotations for 1,773 organismal trait terms, 53,786 functional annotation terms linked to microbes, and 33,305 functional terms are linked to human proteins in the current KG-Microbe graphs.

**Figure 4.**
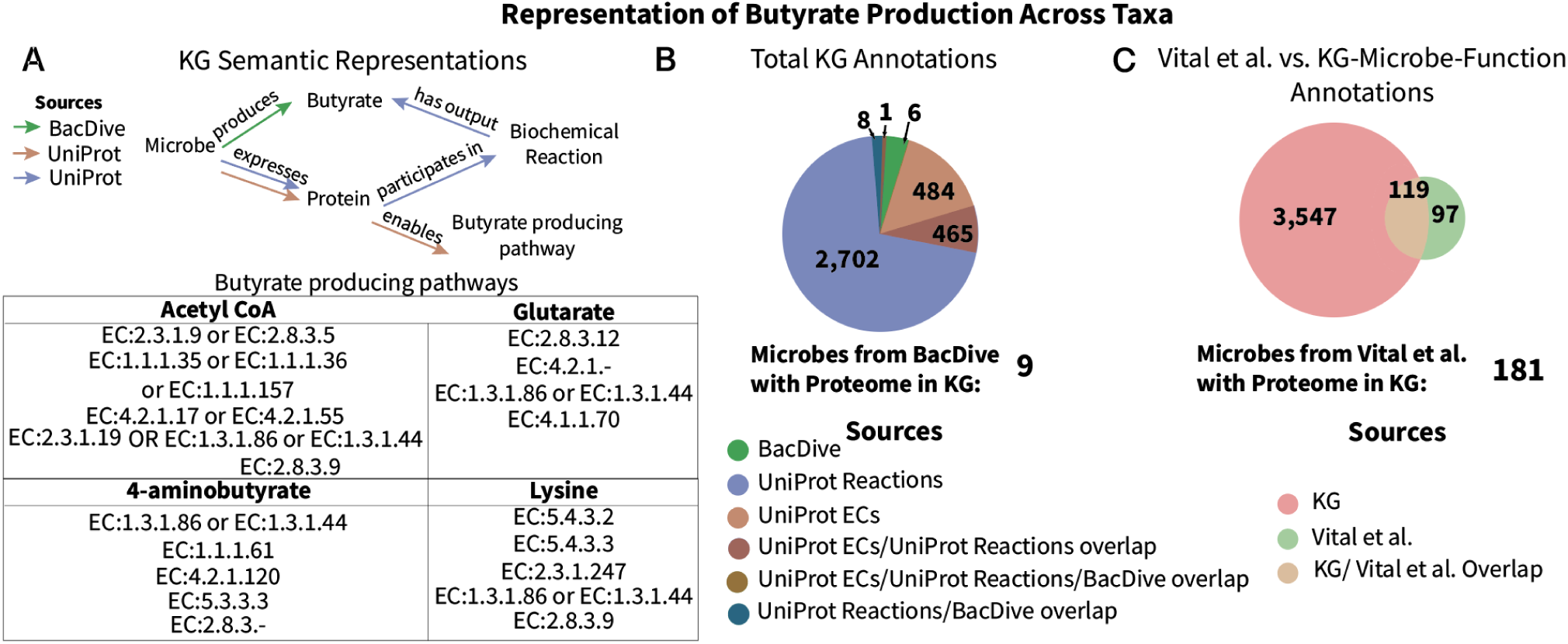
(A) Semantic representation of butyrate production in KG-Microbe-Function. BacDive provides organismal traits specifying which microbes can produce butyrate, and UniProt provides functional annotations through either reactions that have butyrate as a product (predicted using eQuilibrator) or through a curated set of EC numbers that are involved in a butyrate producing pathway. (B) Number of unique microbes that are annotated to produce butyrate based on the semantic representations from (A). The number of taxa from the BacDive set that have UniProt annotations in the KG are also noted. (C) Number of unique microbes that are annotated to produce taxa in Vital et al. (green) or through any representation from (A) (pink). The number of microbes from the Vital et al. set that have UniProt annotations in the KG are also noted.

We compared the set of taxa reported to produce butyrate across these three semantic representations in KG-microbe, as well as those found with a sequence-based approach that serves as an external gold standard of butyrate producers not ingested in the KG. This analysis assesses the correctness of information represented in the KG, where we would expect alignment of the taxa identified through semantic representations to those in the gold standard. We evaluated a pathway prediction analysis performed by Vital et al. In that paper, genomes were analyzed from the Integrated Microbial Genome database to find potential butyrate producers according to EC annotations from KEGG and a hidden Markov model (HMM)-based approach that inferred homologous sequences trained from protein sequences in NCBI of bacteria known to produce butyrate^28,29^. This method predicted 225 microbes capable of butyrate production, of which 216 could be mapped to NCBI Taxonomy identifiers^28^ and 181 had protein annotations from UniProt in KG-Microbe-Function.

A majority of the butyrate producers in the KG were from the semantic representation involving biochemical reactions (3,176 total), followed by the semantic representation involving enzymatic activity (949 total, Figure 3B). Very few direct butyrate production annotations from BacDive exist, suggesting room for an experimentally validated gold standard for this trait (15 total, Figure 3B). All microbes in the BacDive set that had functional annotations in the KG were also expected to produce butyrate based on the reaction or enzymatic activity semantic representation. A Monte Carlo simulation of 1,000 iterations (p < 1e-4) indicated that this overlap was statistically significant and not due to random chance. Thus we show the accuracy of the taxa from UniProt having functional annotations that align with BacDive experimentation.

In comparing the KG butyrate producers to the set in Vital et al., we found that 119 of the 216 taxa (55%) according to Vital et al. were identified using these semantic representations in KG-Microbe-Function. Of the 181 taxa from Vital et al. with functional annotations in the graph set, 117 (65%) were represented in the KG as producing butyrate (the other 2 that overlap with the Vital et al. set were from BacDive, Figure 4C). To understand the predicted set of butyrate producers from KG-Microbe-Biomedical-Function, we examined the taxonomic groups annotated by this trait. We compared taxa that are in the KG and annotated to produce butyrate but not in Vital et al. (3,547 total taxa, Figure 4C) to those that are in Vital et al. but not annotated in the KG (97 total taxa, Figure 4C). A majority of the phyla and families represented in Vital et al. are covered in the KG (Figure 5A) even though there exists 97 taxa from Vital et al. that do not have a KG representation of butyrate production. A majority of the taxa annotated to have this trait by KG-Microbe-Function are also part of the primary phyla present in the human gut: Bacteroidota, Bacillota (Firmicutes), Actinomycetota (Actinobacteria), Pseudomonadota (Proteobacteria), or Fusobacteriota (Fusobacteria), though not all species or strains of that phyla are expected to survive in the human gut (Figure 5A)^2^. These results confirm alignment of taxa at coarser taxonomic resolution (at the phylum level) between the KG and Vital et al. despite having differences at the strain or species level and with KG-Microbe KGs having over am order of magnitude more data. Over time, such discrepancies between prior published results are expected to arise from the continued accumulation of new data, which is captured in frequently updated databases such as UniProt.

**Figure 5.**
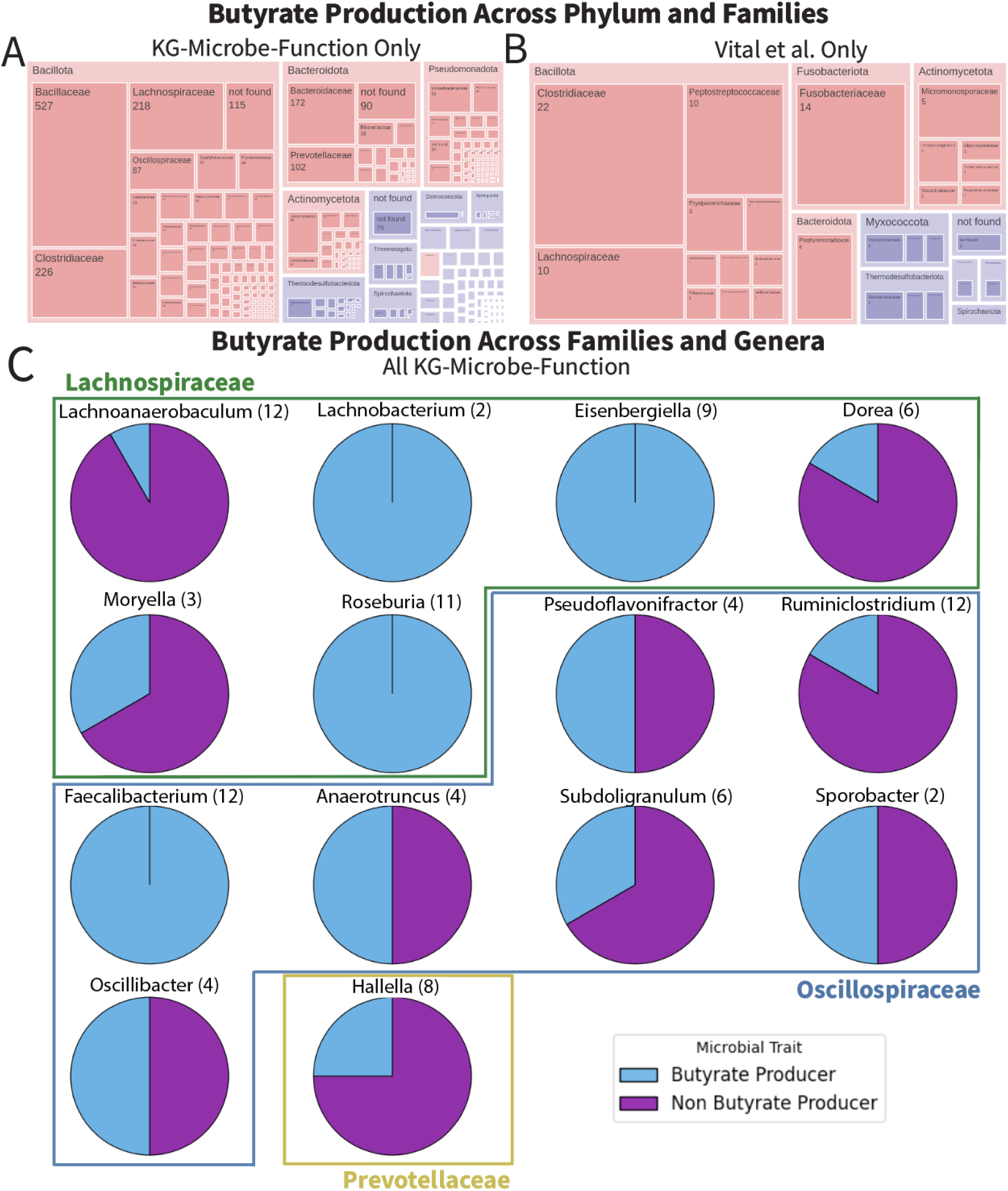
Treemaps of observed phyla and families among (A) annotated microbes in KG-Microbe-Function that are not in Vital et al. set and (B) microbes in the Vital et al. set that are not in KG-Microbe-Function. Red boxes signify the main phyla found to be present in the human gut, while purple boxes signify phyla not commonly found to be present in the human gut. (C) Proportion of butyrate producers among annotated microbes in KG-Microbe-Function, including families understood to be prominent butyrate producers in the gut.

To further assess the accuracy of the KG knowledge, we evaluated the potential for the graph to identify key butyrate producing families in the human gut, including Ruminococcaceae, Lachnospiraceae, Bacteroidaceae, Prevotellaceae, Clostridiaceae, Oscillospiraceae, and Eubacteriaceae^30^. We found that 100% of species and strains in the genera *Faecalibacterium* and *Roseburia* were predicted to be butyrate producers, and 50% of species and strains in the genus *Oscillobacter*.

Through the comparison of semantic representations in the KG, an external gold standard, and what we know about taxa present in the human gut microbiome, we found alignment at the family and phylum levels. We have demonstrated that the large number of taxa represented in the KG though not by the Vital et al. analysis are aligned with our understanding of butyrate production in the gut. The species and strain level differences that arise between Vital et al. and the KG allude to the varying functional annotations, which is expected with manual curation and predictions. This semantic representation methodology thus demonstrates the ability to summarize microbial function according to the modeling of reaction and pathway information alongside phylogenetic relationships within KG-Microbe.

### 2.3. Applications of KG-Microbe in the Context of Disease

We applied the representation of microbial taxonomy and function within KG-Microbe-Biomedical-Function in two disease contexts to demonstrate its usefulness in biomedicine. Examining the role of the microbiome in disease expands our understanding of how microbes can influence human health, either positively or negatively. With population wide studies, microbial function is often predicted at a coarse taxonomic resolution. Through the representation of phylogenetic relationships in the KG, it is possible to explore microbial function at finer taxonomic resolution (the species or strain level), bringing us closer to targeted therapeutics such as probiotics to treat diseases. Butyrate production is a key metabolite for maintaining intestinal barrier function and reducing inflammation, and population studies have found significant reduction in intestinal microbes that produce butyrate in inflammatory bowel disease^26^ and PD^31^, where reduced intestinal barrier function is also observed. In PD, the aggregation of alpha-synuclein, often observed in the enteric nervous system, is heightened by increased inflammation from reduced butyrate production^31^. Population level results are useful to begin to characterize the difference in microbial communities in disease compared to health, but it is difficult to infer functional explanations for these differences due to the coarse taxonomic resolution of the results of 16S rRNA amplicon sequencing.

To explore the functional qualities of these correlative results, we used the taxonomic structure of KG-Microbe-Biomedical-Function to gather the ‘pan-genome’ of all strains below each taxon annotated as increasing or decreasing with disease. We found that a significantly higher number of butyrate producing microbes were associated with a decreased likelihood of both IBD and PD (p = 1e-142, p = 2e-288 respectively) (Figure 6). By traversing the KG in a semantically constrained way we were able to find representative finer-resolved species or strains for each taxon from the original source (labelled as “Original Source Representation” in Figure 6B) and therefore provide a clearer summary of the informative trait of butyrate production.

**Figure 6.**
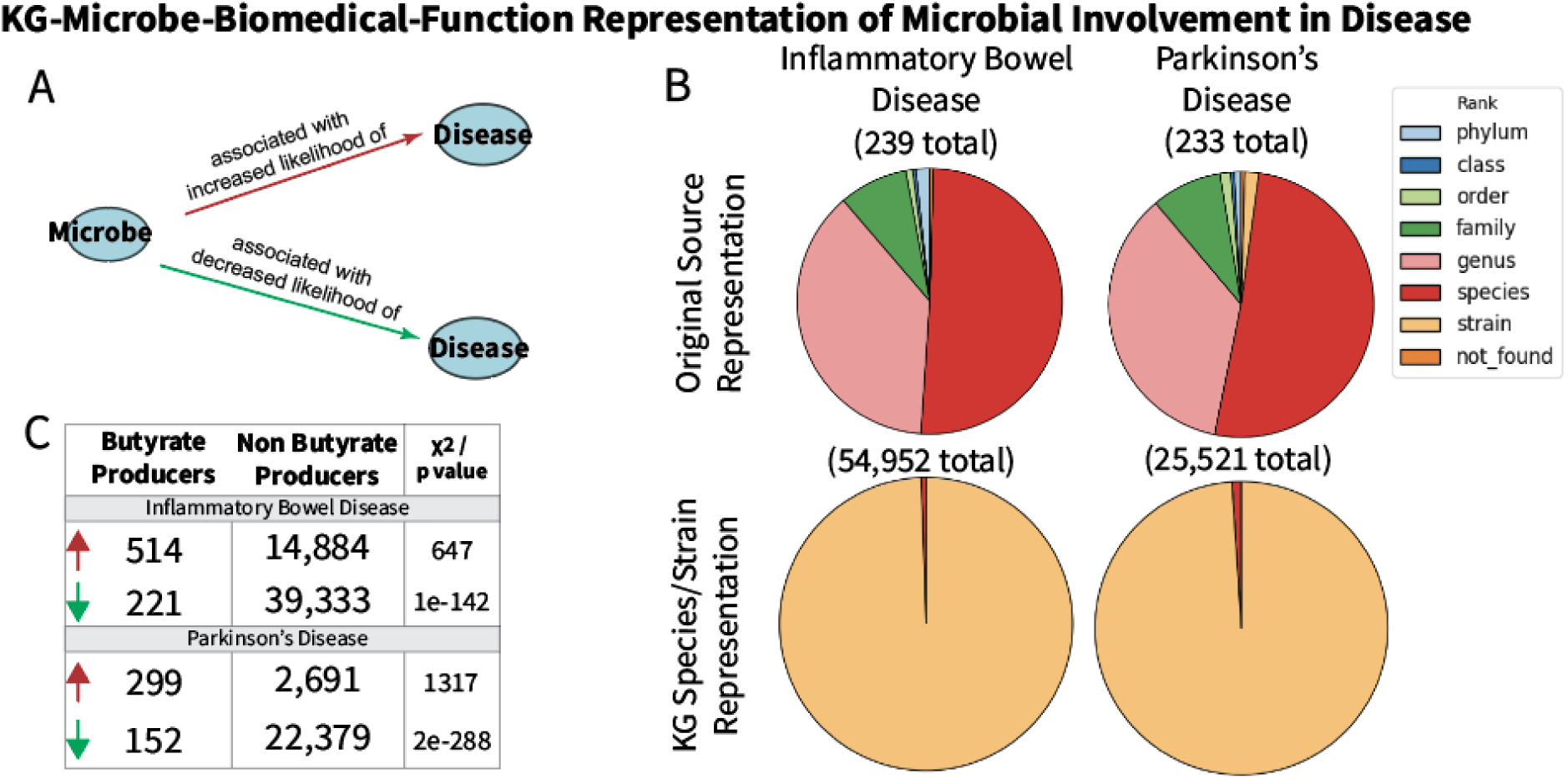
(A) Semantic representation of microbe-disease relationships in KG-Microbe-Biomedical-Function from Disbiome or Wallen et al. (B) The rank of disease annotations found directly in Disbiome or Wallen et al. (top), then transformed by finding all strain or species level children of each taxon (bottom) for both IBD and PD. (C) Using the species/strain representation of disease annotations, the proportion of butyrate producers along with the corresponding chi-squared test statistic comparing taxa annotated to increase (red arrow) or decrease (green arrow) likelihood of disease.

The phylogenetic comparison of butyrate production and disease association analyses are typical examples of general and popular research activities performed with microbial and microbiome data. We have demonstrated the ability of the KG to uncover microbial function at finer taxonomic resolutions than population level results. KG-Microbe enables analysis of such complex data relationships in a semantically- and taxonomically-aware manner while being supportive of the nuances and richness of microbial data.

### 2.4. Environmental Applications of KG-Microbe

Finally, we demonstrate the unique capability of the KG-Microbe framework in an environmental application and as a source of microbial traits for machine learning (ML)-based predictions. While over 10,000 bacterial and archaeal species exist in the human microbiome, the number of species living in Earth’s environments is estimated to be at least in the millions, and with a much larger diversity of environments and physical conditions, most of them inhospitable to humans^32,33^. Due to these differences, applying KG-Microbe to environmental applications utilizes different parts of the KG than when considering only the human microbiome. In addition, the biomedical examples focused on semantic representations and detailed assessments of paths for a specific outcome, whereas in this section we take an untargeted approach and consider all taxa which have an enzyme (EC) or reaction annotation (Rhea). These two node types can be considered equivalent representations in some cases (i.e., the enzyme is semantically equivalent to the reaction it performs), though the outcomes of genome annotation can vary due to sequence similarity cutoffs and differences in manual curation.

Using strains and species with both temperature preference information and genomic annotations of enzymatic activity (as EC nodes) and biochemical reactions (as Rhea nodes), we trained a gradient boosted decision tree classification model to predict four temperature classes: psychrophilic, mesophilic, thermophilic, and hyperthermophilic (see Methods). A semantic representation of genomic annotations was used to produce binary features for taxa (e.g. when the triples *“protein, biolink:participates_in, enzymatic activity”* and *“protein, biolink:derives_from, microbe”* exist, the corresponding *“enzymatic activity”* is included as a feature for that microbe). The model performed well with 90% and 61% precision and recall on withheld test data (macro avg. across four temperature classes). Interpreting the feature importance for hyperthermophilic and psychrophilic taxa (Figure 7) revealed sets of EC and Rhea annotations with opposite importance (positive vs. negative) for these two extreme temperature preferences, corresponding to enzyme and reaction presence/absence patterns.

**Figure 7.**
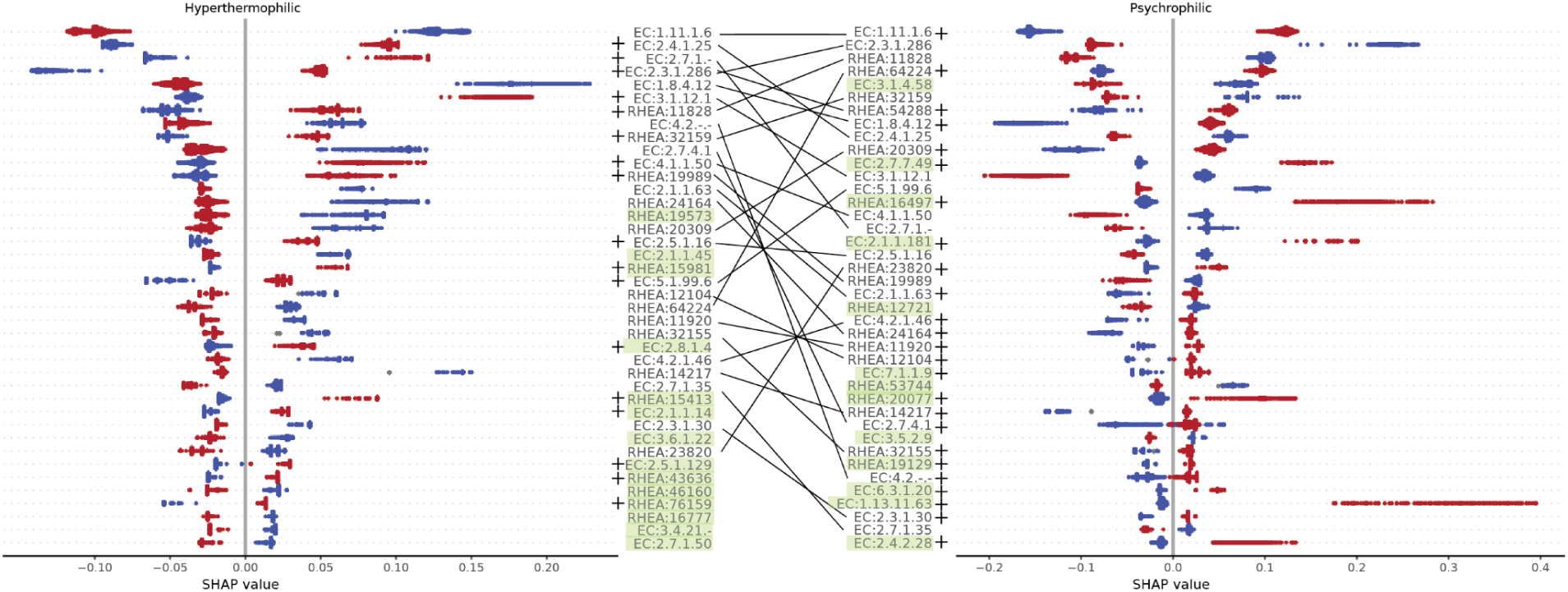
Feature importance for hyperthermophilic (A) and psychrophilic (B) taxa from a multiclass temperature classification model using EC enzyme and Rhea reaction UniProt annotations as features. The top 20 features are shown, sorted in descending order by mean SHAP value. Red corresponds to feature presence and blue corresponds to feature absence. Green background indicates features which are not shared in the top 20 between these two classes, lines connect features with opposite (‘+’ vs ‘-’) importance, and ‘+’ indicates that the presence of that feature has a positive impact on the model.

We considered absolute mean SHapley Additive exPlanations (SHAP) values, which measure the contribution of a given feature to the outcome of a predictive ML model^34^. The top important feature for predicting microbial temperature class preferences was an oxidoreductase acting on peroxide as an acceptor (EC:1.11.1.6), with positive importance for psychrophilic and negative for thermophilic organisms. Oxygen solubility increases with lower temperature and psychrophile metabolism can generate more^35^, hence organisms in these environments could be subject to more oxidative stress. On the other hand hyperthermophiles often inhabit environments with low oxygen levels such as hot springs or deep sea vents and hydrogen peroxide is less stable at high temperatures^36^. The top unique positive feature for hyperthermophiles was adenosylmethionine decarboxylase (Rhea:15981), an enzyme involved in spermidine production, with spermidine known to increase DNA and protein stability^37^. The top unique feature for psychrophiles was RNA-directed DNA polymerase (EC:2.7.7.49), which could be explained by the fact that RNA molecules are highly heat labile and more stable at lower temperatures^38^. In summary, this machine learning application demonstrates that the harmonized knowledge in KG-Microbe can be used to predict broadly relevant biological insights such as temperature preference while providing specific explanations of important microbial features.

## 3. Discussion

The KG-Microbe framework supports the construction of diverse, harmonized, and broadly applicable microbial KGs that support biological discovery. To demonstrate utility and quality of the KGs we performed a series of analyses covering a wide range of applications from biomedical to environmental and beyond. Through evaluation by competency questions, we demonstrated that the harmonization and integration of multiple data sources resulted in an accurate model of microbial genomic traits for applications in mechanistic inferences at varying levels of taxonomic resolution. Incorporating large sources of protein annotations in a semantically consistent structure allowed straightforward inference over the KGs to make predictions about relevant microbial traits such as butyrate production. The applications of this microbial trait analysis in the context of IBD and PD exemplify how the KGs facilitate an understanding of microbial function at finer taxonomic resolutions. This can aid in hypotheses about microbial involvement in disease at the species or strain level, which is important for biomedical applications such as development of biomarkers and targeted therapeutics. The large-scale integration of functional predictions from literature, experimental observation, and predictive models in the KG-Microbe framework offers unrivaled standardization and coverage of microbial traits.

We have also demonstrated the utility of KG-Microbe-Function to support ML-based predictions by transforming or augmenting the knowledge represented into binary features for use in inferences about microbial growth preferences and explanations of the functions involved. This example of collapsing the semantic representation of functional annotations can be extended to other microbial knowledge in the graph, such as organismal traits, or incorporating latent representations of graph structure such as graph embeddings. These applications can be extended to other biological questions to benefit from harmonized knowledge sources.

The four KG-Microbe framework KGs cover a broader scope of knowledge than previously captured in microbial KGs. The MetagenomicKG was recently published as a resource that effectively represents microbial sequences from the Genome Taxonomy Database (GTDB) with microbial functions described in the KEGG database, among other sources, in order to characterize pathogens in relation to human disease phenotypes described in KEGG, the Bacterial and Viral Bioinformatics Resource Center (BV-BRC), or MicroPhenoDB^7,13,39–42^. Because microbes are not mapped to NCBI Taxonomy, the largest taxonomic classification resource, the traversal of phylogenetic relationships among taxa that is possible in all KG-Microbe graphs is not supported by this resource^13^. Other microbial KGs represent a limited number of microbial relationships and contexts. MicrobiomeKG introduces a set of manually curated microbial associations from literature, while MGMLink integrates correlative results from the gutMGene database into a large biomedical KG^8,43^. MiKG4Md and Pre-/Probiotics Knowledge Graph similarly combine data from a series of manuscripts from a directed literature search to represent the role of the microbiome in mental health disorders^9,10^. The KG-Microbe biomedical graphs integrate a wider variety of microbe-disease information, incorporate organismal and genomic traits, and provide more knowledge of host physiology (human in the case of the KGs described) with functional annotations. The KG-Microbe framework fills an important need in building KGs relevant to a variety of microbial contexts that are rooted in broadly relevant knowledge of microbiology.

As with any KG, we are reliant on the knowledge in the ingested sources. UniProt is our main source of functional annotations and was chosen because of its partially reviewed proteome data, broad scope, and standardization of microbial function annotations. In comparison to resources such as the KEGG or MetaCyc, UniProt is freely available, supports frequent updates (as often as monthly), and provides mappings to common primary resources to help with standardization, all of which ensures that the KG-Microbe function graphs are up to date with recent genome annotation knowledge^19,40,44^. A large fraction of the predicted butyrate producing taxa from KG-Microbe-Function are based on UniProt annotations, but further experimentation would be necessary to verify these results. Of the 3,600 taxa predicted to produce butyrate in the KG from the UniProt source, 2,859 are annotated with a “Probable butyrate kinase (BK)”. UniProt relies on both expert and community curation to inform protein annotations, however many of these annotations are not experimentally validated at the function level, and therefore may be misannotated or incomplete. The inclusion of resources like BacDive alongside these functional annotations is important because BacDive organismal traits are derived from literature and in-house laboratory experiments (see Methods) and hence can serve as a near gold standard.

A future direction of this KG framework could be to provide a sequence-integration layer for taxa rather than names mapped to NCBI Taxonomy. By introducing taxa based on sequence integration of whole genomes or by sequence accession identifiers, an additional layer of confidence and provenance is enabled, and this also supports integration of novel taxa not represented in existing taxonomies by linking them to the closest neighbor or appropriate taxonomy rank. Because BacDive provides sequence provenance for each strain-level record, 16S or whole genome sequences can be found from the BacDive source using the corresponding NCBI Taxonomy or BacDive IDs from the KG.

The KGs are also limited by the lack of reaction directionality for genome annotations due to the bi-directional representation of reactions in UniProt. We introduced reaction directions based on the eQuilibrator API, which can be broadly applicable to chemical reaction knowledge represented in this way. A future direction would be to add this to the KG semantic representation of biochemical reactions with additional assumptions (e.g. standard growth conditions). Further, there is potential to incorporate more evidence-based models of microbial metabolism such as Genome-Scale Metabolic Networks (GSMNs). Resources such as the Biochemical, Genetic and Genomic knowledge base (BiGG) or MetaNetX could provide these on a larger scale^45,46^. There are also knowledge bases such as AGORA2, which consolidates GSMNs of gut microbiome-relevant organisms, that can be used in predictive models of microbial biochemical interactions to provide more targeted analyses of specific taxa^47,48^.

The biomedical KGs include microbe-disease correlations from Disbiome, which provides a range of microbe-relevant diseases and correlations from literature. The broad scope of disease from this resource supports KG analyses such as understanding butyrate production in the context of inflammatory diseases (e.g., IBD and PD). However as shown in our disease analysis, Disbiome annotations are at a coarse taxonomic resolution. It would be useful to introduce metagenomics studies of microbes in a disease context to expand the function insight that can be gained from KG-Microbe-Biomedical-Function. Several other structured disease databases exist that could be introduced to the biomedical KG to expand the possibilities of inference in other disease contexts^49^.

The functional KGs constructed with this framework are compliant with the Biolink model, with the exception that they all include potentially redundant node types such as EC, Rhea, and GO, which all represent enzymatic activity in different ways. This supports the complete representation of protein or microbe annotations from all sources (UniProt, BacDive), since not all potentially equivalent annotations are found for each protein. Future versions of the KGs might collapse redundant nodes based on equivalency analysis, for example combining EC and Rhea terms as synonyms which pertain to identical biochemical reactions, to ensure uniqueness in all node types and support more straightforward inference. Combining EC and Rhea nodes can also reduce the number of content edges and provide more straightforward paths describing protein function. These node normalization tasks are important considerations for the modeling choices of each KG built from this framework. Our competency questions demonstrate the importance of fully representing sequence annotations. Furthermore, functional annotation methods for microbes are still being expanded and refined.

The architecture of this framework supports a variety of applications for the resulting KGs, from understanding specific mechanisms of the host microbiome in disease to providing growth preference predictions for microbes. The KG-Microbe framework paves the way for modular KG development, where customizable KGs can be constructed based on standardized transform and modeling patterns for a wide range of data sources. We present targeted use cases for the KGs, with KG-Microbe-Core providing a base for inference and learning about organismal traits of microbes, KG-Microbe-Function enabling the identification of important microbial functions, and KG-Microbe-Biomedical-Function broadening our understanding the host-associated microbiome. With the standardization of both experimental observations and functional predictions according to the Biolink Model, all KG-Microbe KGs demonstrate the ability to draw from the many sources of knowledge available in an accurate way that increases their collective utility.

## 4. Materials and Methods

### 4.1. KG-Microbe Architecture

The download, transform, and merge steps for constructing KGs in the KG-Microbe framework, build upon the KG-Hub construction process and involve three coordinated code repositories: Uniprot2s3, KG-Microbe, and KG-Microbe-Merge (Figure 8)^50^. The uniprot2s3 repository is used to download information using the UniProt API (see the “UniProt” section below) (https://github.com/Knowledge-Graph-Hub/uniprot2s3). The underlying infrastructure that supports the construction is KG-Hub and the KGX tool (version 2.4.2), which provides a template repository and tooling for KG construction^50^. During the transform step, content from all sources is normalized and modeled according to the “KG-Microbe Construction and Modeling” section below by the kg-microbe repository, and redundant edges and nodes are accounted for in cases where overlapping information exists in multiple sources (https://github.com/Knowledge-Graph-Hub/kg-microbe). The resulting transform modules in KGX format are then published in a GitHub release to provide additional data, provenance, and to support interoperability with other sources and KGs. Finally, sources are merged into distinct KG resources. KGX was used to merge KG-Microbe-Core and KG-Microbe-Biomedical in the kg-microbe repository. Due to the size of the microbial UniProt transform, we implemented a merge process using DuckDB to merge the UniProt edges in KG-Microbe-Function and KG-Microbe-Biomedical-Function with other sources in a memory efficient way (https://github.com/Knowledge-Graph-Hub/kg-microbe-merge).

**Figure 8.**
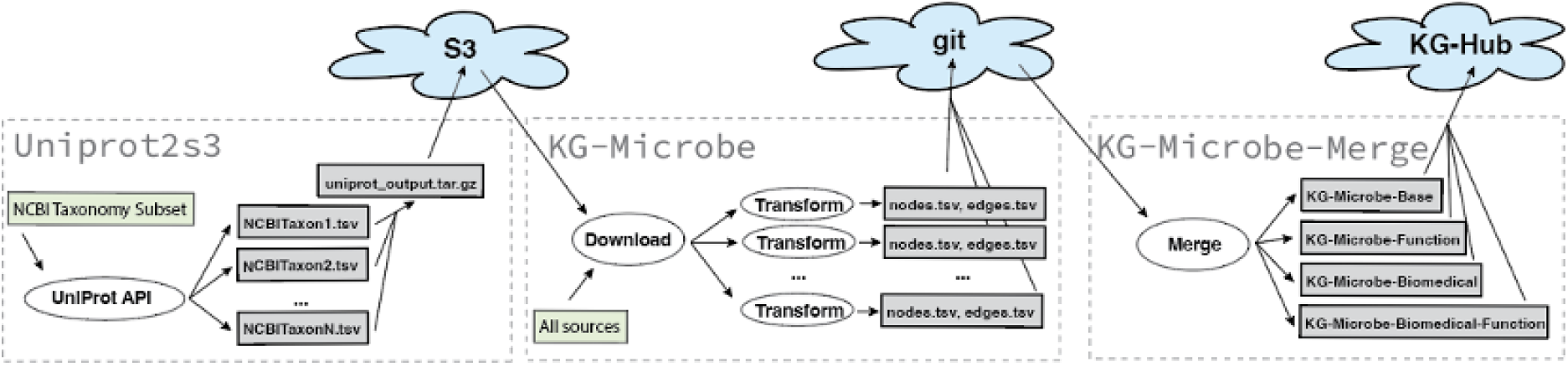
Architecture of the build process for KGs built from the KG-Microbe framework. First Uniprot is queried for all organisms of interest which stores the raw responses in a TSV file per organism. These are compressed into a single data product and stored in S3. Next all input sources are downloaded from S3, the Open Biological and Biomedical Ontology (OBO) Foundry, or cached API downloads in Google Drive as input to the unique transform code for each source. These transforms are then released as compressed files in GitHub or S3 for large files. Last, the transform components are merged according to the desired graph and content. Final products are stored in KG-Hub (KG-Microbe-Core and KG-Microbe-Biomedical) and KG-Registry.

### 4.2. KG Construction and Modeling

The sources ingested for each KG produced by the KG-Microbe framework are shown in Figure 9. The version, data type, and modeling choices of all ingests are detailed in the following sections.

**Figure 9.**
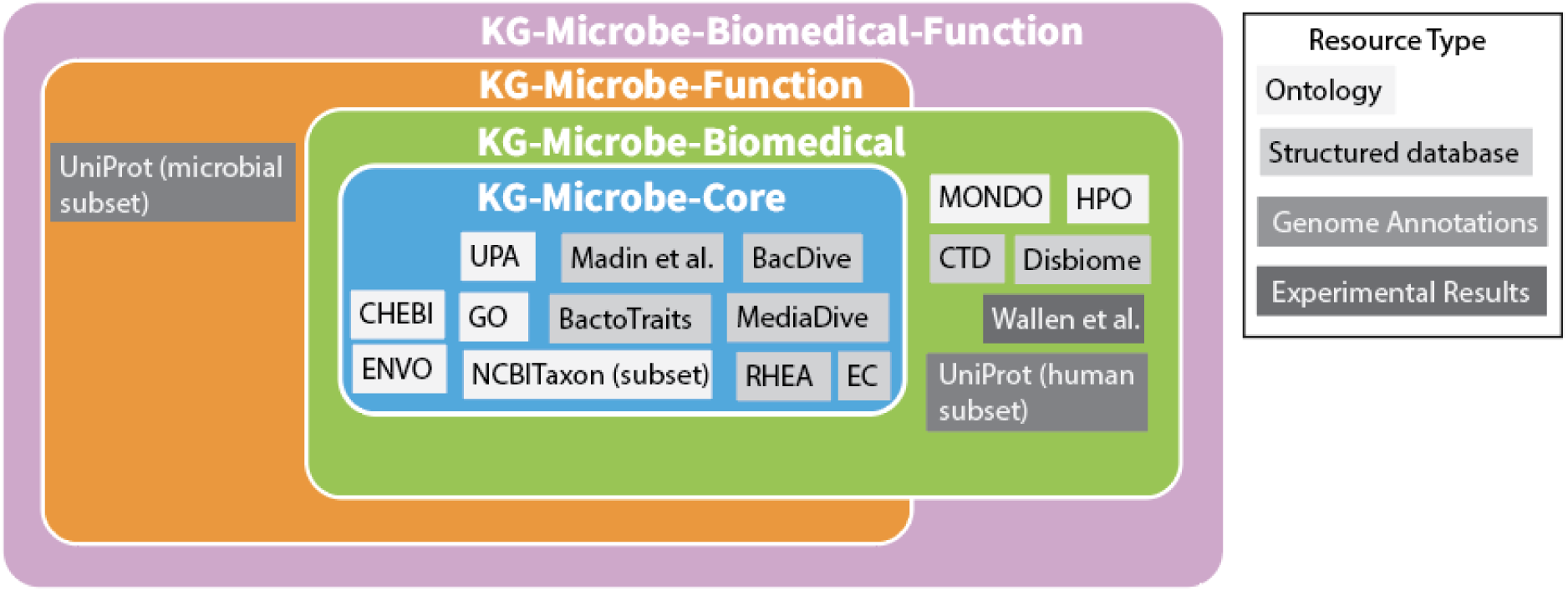
Sources included in each KG generated with the KG-Microbe framework. The type of each source is characterized by box color. Each KG includes ontologies, structured databases, experimental data from the literature, or genome annotations.

#### 4.2.1. BacDive

The BacDive database is included in all KGs (Figure 9). Information from BacDive was obtained using the API and was downloaded on October 3, 2023 as a JSON file. Each strain level taxon has a unique identifier within the JSON file. The strain identifier is the *“NCBI tax id”* subfield of the *“General”* subfield when present, otherwise is given a unique identifier according to the *“BacDive-ID”* field with the custom *“strain”* prefix (e.g., *“strain:bacdive_17187”*). The strain label is formed by combining the *“strain designation”* subfield of *“Name and taxonomic classification”,* the NCBITaxon ID, and the BacDive ID (e.g., “*bacdive_17187 as SH2 of NCBITaxon:1123511”*). The *“Physiology and metabolism”* field is used to find oxygen, salt concentration, metabolism, antibiotic, enzymatic activity, and API or antibiogram assay information. The *“oxygen tolerance”* subfield is used to generate nodes of 9 unique categories: *“oxygen:aerobe”, “oxygen:aerotolerant”, “oxygen:anaerobe”, “oxygen:facultative_aerobe”, “oxygen:facultative_anaerobe”, “oxygen:microaerophile”, “oxygen:microaerotolerant”, “oxygen:obligate_aerobe”,* or *”oxygen:obligate_anaerobe”*. Morphology information is captured from the *“Morphology”* subfield, within which are the *“cell morphology”* and *“pigmentation”* subfields. The *“colony morphology”* and *“multicellular morphology”* subfields were not ingested because they are free text fields. The *“cell morphology”* specifies the *“gram stain”* trait (*“negative”* or *“positive”*) which results in three unique nodes: “*gram_stain:positive”, “gram_stain:negative”, “gram_stain:variable”*. 18 unique values from the *“cell shape”* subfield were introduced to the graph with the *“biolink:has_phenotype”* predicate. Whether a microbe is motile or non-motile is sourced from the*“motility”* subfield, introducing the nodes *“motility:motile”* and *“motility:non_motile”.* Temperature categories are extracted from the *“culture temp”* subfield, where only the *“range”* subfield was used to obtain one of four categories (*“temperature:hyperthermophilic”, “temperature:mesophilic”, “temperature:psychrophilic”,* or *“temperature:thermophilic”*), and the quantitative measurements from the *“temperature”* subfield were not used. Salinity tolerance was introduced from the *“halophily_level”* subfield of the *“halophily”* subfield as 7 unique categories (“*extremely_halophilic”, “haloalkaliphilic”, “halophilic”, “halotolerant”, “moderately_halophilic”, “non_halophilic”, “slightly_halophilic”*). Chemical production or consumption results are obtained from the *“metabolite utilization”* subfield, where the associated ID comes from the *“Chebi-ID”* subfield when present, the *“biolink:produces”* predicate is used when a *“+”* is present in the *“utilization activity”* field, and the *“biolink:consumes”* predicate is used when a *“-”* is present in the *“utilization activity”* field. Chemical terms are mapped using provided mappings or by manual curation for the case of 9 terms (using the *“metabolite_mapping.json”* file). When no matching ChEBI exists, chemicals are represented with CAS-RN or KEGG identifiers according to BacDive (Table 1). Chemical production results are also found from the *“metabolite production”* subfield, where *“production”* is *“yes”*. Enzymatic activity is found in the *“enzymes”* subfield and modeled using the *“biolink:capable_of”* predicate from the microbe to the enzyme in the *“ec”* subfield when *“activity”* is *“+”*. Negative results from the *“enzymes”* subfield are not included in the graph (e.g. when *“activity”* is *“-”*). A microbe is considered sensitive to a given chemical when the *“is sensitive”* subfield within *“antibiotic resistance”* is *“yes”*, and resistant when *“is resistant”* is *“yes”*. The negative results of *“is sensitive”* or *“is resistant”* as *“no”* were not ingested. The sensitivity concentration from the “*sensitivity conc.”* subfield is not ingested but is available in the raw BacDive source files. Antibiotic resistance and sensitivity information is also taken from the antibiogram results in the *“antibiogram”* subfield. This field provides a growth medium specified in the *“medium”* subfield and a range of chemical subfields with an associated value specifying the zone of inhibition (ZOI) diameter in millimeters, indicating the extent to which the chemical (an antibiotic) inhibits growth of the strain. Each individual antibiogram result is mapped to a standardized chemical identifier (using the *“metabolite_mapping.json”* file), and the value is returned as a single floating-point number where a range is represented by the mean (e.g. *“42-44”* becomes *“43”*) and a greater than inequality is represented by the corresponding minimum value (e.g. *“>50”* becomes *“50”*). Strains with ZOI values of less than 15mm for a given chemical are considered resistant to that antibiotic, and values of greater than 25mm are considered as sensitive to that chemical; data with values between 15 and 25mm are discarded. Microbes are linked to a chemical via the *“biolink:associated_with_sensitivity_to”* predicate when sensitive and the *“biolink:associated_with_resistant_to”* predicate when resistant. The *“metabolite tests”* and *“fatty acid profile”* subfields are not ingested. The *“Physiology and metabolism”* field is also used to find trophic type information, from the *“nutrition_type”* subfield, resulting in 28 unique categories from the *“type”* subfield (e.g., *“trophic_type:autotrophy”, “trophic_type:chemotrophy”, “trophic_type:lithoautotrophy”*). The *“spore_forming”* subfield was used to find whether or not a microbe can form spores based on the *“yes”* or *“no”* result (*“sporulation:spore_forming”* and *“sporulation:non_spore_forming”*).

**Table 1.** Description of all resources included in the KG-Microbe framework.

| Node Type(s) Represented | Standardized Mappings |
| --- | --- |
| Organism<sup>40</sup> | NCBI Taxonomy (NCBITaxon)<sup>13</sup>, Custom “*strain*” (Bacterial Diversity Metadatabase (BacDive)<sup>14</sup>) |
| Chemical | Chemical Entities of Biological Interest (ChEBI)<sup>62</sup>, CAS Registry (CAS-RN)<sup>63</sup>, PubChem<sup>64</sup>, Kyoto Encyclopedia of Genes and Genomes (KEGG) |
| Antibiotic | Chemical Entities of Biological Interest (ChEBI)<sup>62</sup> |
| Isolation Source | Chemical Entities of Biological Interest (ChEBI)<sup>62</sup>, Environment Ontology (ENVO)<sup>54</sup>, Plant Ontology (PO)<sup>55</sup>, Phenotype and Trait Ontology (PATO)<sup>58</sup>, Uberon Ontology (UBERON), Custom “*isolation source*” (Bacterial Diversity Metadatabase |
|  | (BacDive)<sup>14</sup> and Madin et al.<sup>16</sup>) |
| Enzymatic Activity | Enzyme Commission (EC)<sup>65</sup> |
| Optimal pH | Custom “*pH_opt*” (BactoTraits<sup>15</sup>) |
| pH Range | Custom “*pH_range*” (BactoTraits<sup>15</sup>) |
| pH Spread | Custom “*pH_delta*” (BactoTraits<sup>15</sup>) |
| Temperature Category | Custom “*temperature*” (BactoTraits<sup>15</sup>) |
| Optimal Temperature | Custom “*temp_opt*” (BactoTraits<sup>15</sup>) |
| Temperature Range | Custom “*temp_range*” (BactoTraits<sup>15</sup>) |
| Temperature Spread | Custom “*temp_delta*” (BactoTraits<sup>15</sup>) |
| Optimal NaCl Concentration | Custom “*NaCl_opt*” (BactoTraits<sup>15</sup>) |
| NaCl Concentration Range | Custom “*NaCl_range*” (BactoTraits<sup>15</sup>) |
| NaCl Concentration Spread | Custom “*NaCl_delta*” (BactoTraits<sup>15</sup>) |
| Oxygen Category | Custom “*oxygen*” (Bacterial Diversity Metadatabase (BacDive)<sup>14</sup>, BactoTraits<sup>14</sup>, Madin et al.<sup>16</sup>) |
| GC Content | Custom “*gc*” (BactoTraits<sup>15</sup>) |
| Motility | Custom “*motility*” (Bacterial Diversity Metadatabase (BacDive)<sup>14</sup>, BactoTraits<sup>15</sup>) |
| Trophic Type | Custom “*trophic_type*” (Bacterial Diversity Metadatabase (BacDive)<sup>14</sup>, BactoTraits<sup>15</sup>, Madin et al.<sup>16</sup>) |
| Cell Width | Custom “*cell_width*” (BactoTraits<sup>15</sup>) |
| Cell Length | Custom “*cell_length*” (BactoTraits<sup>15</sup>) |
| Cell Shape | Custom “*cell_shape*” (Bacterial Diversity Metadatabase (BacDive)<sup>14</sup>, BactoTraits<sup>15</sup>, Madin et al.<sup>16</sup>) |
| Sporulation | Custom “*sporulation*” (Bacterial Diversity Metadatabase (BacDive)<sup>14</sup>, BactoTraits<sup>15</sup>) |
| Pathway | Custom “*pathways*” (Madin et al.<sup>16</sup>) |
| Biosafety Level | Custom “*BSL*” (Bacterial Diversity Metadatabase (BacDive)<sup>14</sup>) |
| Medium | Custom “*medium*” (Bacterial Diversity Metadatabase (BacDive)<sup>14</sup>, MediaDive<sup>18</sup>) |
| Solution | Custom “*solution*” (MediaDive<sup>18</sup>) |
| Ingredient | Chemical Entities of Biological Interest (ChEBI)<sup>62</sup>, CAS Registry (CAS-RN)<sup>63</sup>, PubChem<sup>64</sup>, Kyoto Encyclopedia of Genes and Genomes (KEGG)<sup>40</sup>, Custom “*ingredient*” (MediaDive<sup>18</sup>) |
| Assay | Custom “*assay*” (Bacterial Diversity Metadatabase (BacDive)<sup>14</sup>) |
| Protein | Uniprot Knowledgebase (UniProt)<sup>19</sup> |
| Biological process | Gene Ontology (GO)<sup>66</sup> |
| Molecular function | Gene Ontology (GO)<sup>66</sup> |
| Cellular Component | Gene Ontology (GO)<sup>66</sup> |
| Disease | Mondo Disease Ontology (MONDO)<sup>5</sup>, Human Phenotype Ontology (HPO)<sup>67</sup>, Gene Ontology (GO)<sup>66</sup> |
| Human Phenotype | Human Phenotype Ontology (HPO)<sup>67</sup> |
| Metabolic pathway | UniPathways (UPA)<sup>20</sup> |
| Human Gene | HUGO Gene Nomenclature Committee (HGNC)<sup>68</sup> |
| Biochemical Reaction | Rhea<sup>69</sup>, “*UPA:UCR*” (Unipathways Ontology (UPA)) |
| Metabolic Pathway | “*UPA:UPA*” (Unipathways Ontology (UPA)) |

Isolation source information was sourced from the *“isolation source categories”* subfield within the *“Isolation, sampling and environmental information”* subfield. BacDive models isolation sources in categories from 1 to 3, and all three categories were included as new nodes in the graph with the *“isolation_source”* prefix (e.g. *“isolation_source:environmental”*). These standardized values from BacDive were not mapped to any ontology (Table 1).

Biosafety level categories of 1, 2, or 3 were sourced from the *“Safety Information”/”risk assessment”* subfield. The *“biosafety level”* subfield was annotated with the corresponding safety level, which was introduced as a new node (e.g. *“BSL:1”)* and linked to the microbe via the *“biolink:associated_with”* predicate.

Culture medium information is found in the *“Culture and growth conditions”* field, where each medium is assigned a unique integer according to the MediaDive resources (e.g., *“medium:1”*). When the *“growth”* subfield is *“yes”*, an edge is added between the associated organism (e.g., *“NCBITaxon:1396, biolink:occurs_in, medium:1”*). Medium ingredients are ingested from MediaDive (see the MediaDive section of the Methods).

Each API assay in BacDive assesses a number of enzymatic activities, and the combination of the API assay kit type (according to the name of a subfield starting with *“API”*), and the enzymatic activity, when annotated with a *“+”*, was used for each assay node. For example, the *“API zym”* field had 20 unique enzymatic activity results for a given taxon, one of which was being positive for *“Alkaline phosphatase”*, therefore a new node *“assay:API_zym_Acid phosphatase”* was linked to the associated microbe via the *“biolink:assesses”* predicate. Negative results were not included (e.g. when a given enzymatic activity was *“-”*). The antibiotic results are not consistent with the Clinical and Laboratory Standards Institute (CLSI) and European Committee on Antimicrobial Susceptibility Testing (EUCAST) and therefore not for clinical use^51^.

Other fields that were not used from the BacDive JSON file were: *“Sequence information”, “External links”, “Publications”, “Reference”*, *“Sequence Information”, “Isolation”, “Literature”, “Fatty Acid Profile”, “Compound Production”, “Tolerance”, “Culture Temp.”, “Halophily”,* and *“Culture pH”* because they were in free text form.

#### 4.2.2. Madin et al

The Madin et al. ingest is included in all KGs (Figure 9). Phenotype data is sourced from the *condensed_traits_NCBI.csv* from the Madin et al. GitHub repository^52^. Traits were introduced for each organism from the *“tax_id”* column which contained a NCBI Taxonomy ID. The *“metabolism”* column is used to extract information about oxygen tolerance, resulting in six unique categories (*”oxygen:aerobe”, “oxygen:anaerobe”, “oxygen:facultative_anaerobe”, “oxygen:microaerophile”, “oxygen:obligate_aerobe”,* or *”oxygen:obligate_anaerobe”)*. The two unique annotations of *“strictly anaerobic”* and *“anaerobic”* were collapsed into one KG node *“oxygen:anaerobe”*. The *“pathways”* column sources 100 unique pathways that are linked to the corresponding microbe with the *“biolink:capable_of”* predicate. No pathways are mapped to ontologies or external references but instead have the *“pathways”* prefix (e.g. *“pathways:thiosulfate_oxidation”*) (Table 1). The *“carbon_substrates”* column is used to identify chemicals that the corresponding taxa consumes, where all chemicals are mapped to a ChEBI identifier and linked to the microbe with the *“biolink:consumes”* predicate using named entity recognition and manual mappings for 6 unique terms (found in the *“chebi_manual_annotation.tsv”* file) (Table 1). 19 unique cell shapes are sourced from the *“cell_shape”* column (e.g. *“cell_shape:bacillus”*) and linked to the corresponding microbe with the *“biolink:has_phenotype”* predicate. Finally, environments from which microbes are found is sourced from the *“isolation_source”* column, which results in five unique categories that were not mapped to a standardized identifier (*“isolation_source:food_fermented”, “isolation_source:host_animal_ectotherm”, “isolation_source:host_animal_endotherm”, “isolation_source:host_animal_endotherm_intratissue”, “isolation_source:host_animal_endotherm_rumen”*), and the rest of which were mapped to the Environment Ontology (ENVO), ChEBI, the Plant Ontology (PO), the Food Ontology (FOODON), the Phenotype and Trait Ontology (PATO), or the Uberon Ontology (UBERON) using the *data/conversion_tables/environments.csv* from Madin et al. (Table 1)^53–58^.

#### 4.2.3. BactoTraits

The BactoTraits database is included in all KGs (Figure 9). The BactoTraits database was downloaded as a tsv file (*BactoTraits_databaseV2_Jun2022.csv*). BactoTraits provides organismal traits for the same set of taxa at the strain or species level available in BacDive. All columns were mapped to unique prefix and categories using a LinkML structured yaml file called the *“custom_curies.yaml”*. Bacterial size information is binned into four qualitative labels of cell length and width (*very_low, low, mid,* or *high*) each represented as a node. Bacterial cells of length <= 1.3um are connected to the *“cell_length:very_low”* node, those of length 1.3um to 2um to the *“cell_length:low”* node, those of length 2um to 3um to the *“cell_length:mid”* node, and those of length >= 3um to the *“cell_length:high”* node. Similarly, bacterial cell width is represented in ranges of <=0.5um (“*cell_width:very_low”),* 0.5um to 0.65um (“*cell_width:low”),* 0.65um to 0.9um (“*cell_width:mid”),* >=0.9um (“*cell_width:high”)*. 9 unique cell shapes are sourced from BactoTraits. When a microbe was linked to multiple shapes in BactoTraits, KG-Microbe represents each trait as its own node (e.g. “*S_star_dumbell_pleomorphic*” in BactoTraits becomes *“cell_shape:star_dumbbell_pleomorphic”* and the individual linked nodes *“cell_shape:star”, “cell_shape:dumbell”, “cell_shape:pleomorphic”* in KG-Microbe). Growth conditions for each microbe were introduced from BactoTraits, including the optimal range for growth, the range in which growth is possible, and the ecological valence, or spread, of the range in which growth can occur. All of these ranges were binned into unique qualitative labels according to the representations of categories in BactoTraits (very low, low, mid1, mid2, mid3, mid4 and high). For example, salt concentration range nodes vary from *“NaCl_range:very_low”* (≤1% NaCl in BactoTraits) to *“NaCl_range:high”* (>8% NaCl in BactoTraits), whereas the spread of temperature range nodes vary from *“temp_opt:very_low”* (≤10°C) to *“temp_opt:high”* (>40°C), with intermediate ranges: *“temp_opt:low”* (10–22°C), *“temp_opt:mid1”* (22–27°C), *“temp_opt:mid2”* (27–30°C), *“temp_opt:mid3”* (30–34°C), and *“temp_opt:mid4”* (34–40°C). Trophic types were ingested as 10 categories directly from BactoTraits (e.g. *“trophic_type:autotrophy”*, *“trophic_type:chemotrophy”*, *“trophic_type:copiotrophy”*). Gram staining phenotypes were ingested directly from BactoTraits as *“gram_stain:positive”, “gram_stain:negative”, “gram_stain:variable”*, as was sporulation information as *“sporulation:spore_forming”* and *“sporulation:non_spore_forming”* and motility as *“motility:motile”* or *“motility:non_motile”.* The pigmentation of the microbial cell is recorded as 10 unique pigments (e.g, *“pigment:pink”, “pigment:black”, “pigment:carotenoid”*) that are not mapped to an ontology.

#### 4.2.4. MediaDive

The MediaDive database is included in all KGs (Figure 9).MediaDive was downloaded from the REST API v0.1.0 (https://mediadive.dsmz.de/rest/media). All unique media is taken from the *“id”* subfields of the *“data”* field. The DSMZ growth medium IDs that are introduced from BacDive correspond to those in MediaDive, where the *“name”* subfield serves as the node label for each medium. The medium type is extracted from the *“complex_medium”* subfield, which is a *“1”* if complex and *“0”* if not complex, resulting in two medium types: *“medium-type:complex”* and *“medium-type:defined”*. Given each unique medium ID, data from the MediaDive REST API is then used to find the strains that grow in the media using the *“medium-strains”* field, *“bacdive_id”* subfield. The BacDive ID is linked to the NCBI Taxonomy or unique strain node provided by BacDive using the *“biolink:occurs_in”* predicate. The *“medium”* field along with each unique medium ID is used to extract information about the medium content via the *“solutions”* subfield, where a new node is introduced using the *“id”* subfield (e.g. *“solution:1”*), and all ingredients are found in the *“recipe”* subfield using *“compound_id”*. The *“ingredient”* field along with each unique compound ID is used to represent each chemical as ChEBI, KEGG, PubChem, or CAS-RN IDs (using the *“ChEBI”, “KEGG-Compound”, “PubChem”,* or *“CAS-RN”* subfields respectively) (Table 1). When no chemical mapping was available, the custom *“ingredient”* prefix was used. Two new triples of the form *“solution, biolink:has_part, chemical”* and *“medium, biolink:has_part, solution”* are added using this content.

#### 4.2.5. Rhea

The Rhea database is included in all KGs (Figure 9). Biochemical reaction knowledge from the Rhea database is retrieved from Expasy flat files. The PyOBO Python package enables the construction of OBOs from structured databases and is used to retrieve relationships among biochemical reactions, and between biochemical reactions and chemicals. As part of PyOBO, the *“rhea-directions.tsv”* and *“rhea-relationships.tsv”* files are used to model the direction of each biochemical reaction and the hierarchy of biochemical reactions, respectively. Rhea terms have IDs in multiples of 4 starting at 10000, where the first is *“undirected”*, the second is *“left to right”*, the third is *“right to left”*, and the fourth is *“bidirectional”*. Each unique Rhea term is related to its parent term (except for the *“undirected”* term) via the *“biolink:subclass_of”* predicate. This pattern is used to model the input and output chemicals accordingly. The *“biolink:input_of”* and *“biolink:output_of”* predicates are used to link chemicals to reactions assigned with *“left to right”* or *“right to left”* directions. The *“biolink:participant_of”* predicate is used to link chemicals to reactions that are *“undirected”* or *“bidirectional”*. In addition to using the Rhea model from PyOBO, the *“rhea2ec.tsv”* and *“rhea2go.tsv”* files are downloaded as sources for relationships between biochemical reactions and molecular functions (modeled in the form *“molecular function, biolink:enables, biochemical reaction)*, and relationships between biochemical reactions and enzymatic activity (modeled in the form *“biochemical reaction, biolink:enabled_by, enzymatic activity*).

#### 4.2.6. Ontologies

The Mondo Disease Ontology (MONDO) and human phenotype ontology (HPO) are only included in KG-Microbe-Biomedical and KG-Microbe-Biomedical-Function, while all other ontologies are included in KG-Microbe-Core and KG-Microbe-Function as well (Figure 9). Ontologies are downloaded as JSON or OWL files from the Open Biological and Biomedical Ontology (OBO) Foundry^50^. GO, HPO, and ENVO are transformed into edges and nodes using the *transform* function from KGX (version 2.4.2). EC, UPA, ChEBI, and MONDO are first transformed via the KGX library followed by post processing steps to correctly model the data for the purposes of KG-Microbe. In this step, the following prefixes are replaced from the original file: *“eccode”* with *“EC”* (from the EC JSON file), *“CAS”* with *“CAS-RN”* (from the ChEBI OWL file), *“OBO:UPa_UCR*” and *“OBO:UPa_UPA”* with *“UPA”* (from the UPA OWL file), and *“*http://identifiers.org/hgnc/*”* with *“HGNC”* (from the MONDO JSON file). UPA chemical reactions are mapped to Rhea terms using the *“xref”* field when possible, otherwise they keep the given ID with the *“UPA:UCR”* prefix (e.g. *“UPA:UCR00014”*). We derived the transitive relationships from UPA for triples of the form *“enzymatic reaction, part of, linear sub pathway”* and *“linear sub pathway, part of, pathway”*. In this process, enzymatic reactions were mapped to GO terms while pathways were prefixed with *“UPA:UPA”*, resulting in new triples of the form *“molecular function, biolink:enabled_by, metabolic pathway”* and *“biological process, biolink:related_to, metabolic pathway”*. Similarly, we derived transitive relationships for triples of the form *“chemical reaction, part of, enzymatic reaction”*, *“enzymatic reaction, part of, linear sub pathway*”, and *“linear sub pathway, part of, pathway”*. Here, new triples of the form *“biochemical reaction, biolink:part_of, metabolic pathway”* were introduced to the KG.

As part of the KG-Microbe construction we generated a trimmed version of the NCBI Taxonomy that only included the single cell organisms of interest. The following eukaryotic or otherwise irrelevant taxonomic parent nodes and their descendants were removed: other sequences (NCBITaxon:28384), Viridiplantae (NCBITaxon:33090), Metazoa (NCBITaxon:33208), herbal medicine (NCBITaxon:1407750), mixed DNA libraries (NCBITaxon:1145094), mixed EST libraries (NCBITaxon:119167), mixed libraries (NCBITaxon:590738), Viruses (NCBITaxon:10239). Trimming was performed using ROBOT (ROBOT is an OBO Tool), a command line tool that supports filtering of OBOs in a logically consistent manner among other operations supporting ontology development^59^.

#### 4.2.7. UniProt

The microbial UniProt ingest is included in KG-Microbe-Function and KG-Microbe-Biomedical-Function (Figure 9). Protein annotation data for KG-Microbe is sourced from UniProt proteomes, which are sets of proteins linked to a given taxa^19^. The KG-Microbe pipeline extracts functional annotations from the UniProt REST API using parallelization with multiple CPU cores and produces a flat file for each NCBITaxon organism. Query fields used to construct the KG for both the microbial subset and humans are *“xref_proteomes”*, *“organism_id”*, *“accession*”, *“protein_names”*, *“ec”*, *“ft_binding”*, “*go”*, *“rhea”*, and *“cc_function”*. Information about whether a given protein is reviewed or unreviewed is ingested directly from UniProt, though not included in the graphs. The infrastructure is supported in the uniprot2s3 GitHub repository, which outputs all organism specific files and stores them as a compressed file in S3. This serves as the raw protein functional annotation source data endpoint used to produce the transformed annotation data for the merged KGs (Figure 7). Provenance of whether a protein is from SwissProt or TrEMBL is recorded in the *“reviewed”* field of the raw data, though not included in the final graph. The resulting UniProt raw files and other resources are then downloaded to prepare for extracting and modeling the information (the transform step in Figure 7). The *“organism_id”* field corresponds to an NCBI Taxonomy ID to which all functional information is linked. The *“accession”* field is used for each unique protein ID (e.g. *“UniprotKB:A0A1H0WGA5”)*, which is linked to the microbe via the *“biolink:derives_from”* predicate. The protein label is extracted from the *“protein_names”* field, where only names are included (in some cases more than one, semicolon delimited) and any EC IDs are excluded. Biological process, molecular function, and cellular component GO terms are extracted from the *“go”* field, excluding any text or brackets, and linked to microbes via the predicate “*biolink:participates_in”, “biolink:participates_in”,* or *“biolink:located_in”*, respectively. The GO term categories (GO-BP, GO-MF, GO-CC) are differentiated using the Ontology Access Kit (OAK)^60^. Biochemical reactions are extracted as Rhea IDs from the *“rhea”* field and linked to organisms via the *“biolink:participates_in”* predicate because all Rhea terms are undirected, and enzymatic activities as EC IDs from the *“ec”* field are linked to microbes via the *“biolink:enables”* predicate. Only ChEBI terms are extracted from the content in the *“ft_binding”* field to yield chemicals to which the corresponding protein binds via the *“biolink:binds”* predicate.

The human UniProt ingest is included in KG-Microbe-Biomedical and KG-Microbe-Biomedical-Function (Figure 9). For the human query only, the *“gene_primary”* and *“cc_disease”* fields are also included. The *“gene_primary”* field contains a gene name which is mapped to a HUGO Gene Nomenclature Committee (HGNC) ID using MONDO in order to align with MONDO genes. The *“cc_disease”* field contains Online Mendelian Inheritance in Man (OMIM) IDs for diseases associated with the corresponding protein. These terms are mapped to MONDO terms using the *“xref”* fields of the MONDO OWL file. The protein is linked to the disease via the *“biolink:contributes_to”* predicate. We obtained UniProt proteome annotation data for 29,154 taxa represented by 56,670 proteomes including 5,739 plasmids. Organisms included in the UniProt ingest are limited to those associated with a proteome (reference or non-reference proteome) and over 1,000 total proteins across all linked proteomes (Supplementary Figure 1). Only the reference proteome was included for the human set of proteins (*“UP000005640”*), as well as unreviewed proteins. This results in potentially redundant human proteins which ensure that no annotation information is lost. This protein number cutoff was chosen based on computational constraints, such that parallel processing allowed all entries to be processed within the transform step on a single machine with 512GB of RAM. The human subset of the UniProt database is linked to the corresponding NCBITaxon ID (NCBITaxon:9606) though it is not linked to the taxonomic hierarchy in the graphs.

#### 4.2.8. Disbiome

The Disbiome ingest is included in the KG-Microbe-Biomedical and KG-Microbe-Biomedical-Function graphs (Figure 9). Disbiome relationships were downloaded from the Disbiome API as a JSON file. The disease name was extracted from the *“disease_name”* field. The disease was normalized by a string match to MONDO, HPO, or GO terms, or by manual curation if no match was found automatically (Table 1). The organism name was extracted from the *“organism_name”* field and the organism ID from the *“organism_id”* field, which corresponded to an NCBI Taxonomy ID when present. To map microbes to NCBI Taxonomy terms when no organism ID was present, a direct string match between the microbe name in Disbiome and that in NCBI Taxonomy was first attempted. If no matches were found, brackets were included around the first term, assumed to be a genus name, and the match was attempted again. For example, after no NCBI Taxonomy match was found for “*Ruminococcus lactaris”*, “[*Ruminococcus] lactaris”* was successfully mapped to “*NCBITaxon:46228”*. Any remaining microbes were then manually mapped to a given NCBI Taxonomy. The predicate between a microbe and disease was determined from the *“qualitative_outcome”* field, where the predicate *“biolink:associated_with_increased_likelihood_of”* was used for the *“Elevated”* outcome and *“biolink:associated_with_decreased_likelihood_of”* was used for the *“Reduced”* outcome.

#### 4.2.9. Wallen et al

Data from *“Supplementary Table 1”* from Wallen et al. was downloaded for extraction of relationships between microbes and PD. Microbe names were extracted from the *“Species”* column. Similar to Disbiome, a direct string match between the microbe name in Wallen et al. and that in NCBI Taxonomy was first attempted, followed by the microbe name with brackets around the first string, followed by manual mapping. PD was manually assigned the MONDO term *“MONDO:0005180”*. The direction of the relationship between a given microbe and PD was determined based on the *“RA in PD”* and *“RA in NHC”* columns, which specify the relative abundance of the microbe in the disease and control groups, respectively, using Metagenomic Phylogenetic Analysis (MetaPhlAn). The *“biolink:associated_with_increased_likelihood_of”* predicate was used when the abundance in the disease group was greater than in the control group, while the *“biolink:associated_with_decreased_likelihood_of”* predicate was used when the abundance in the control group was greater than in the disease group. Only significantly differentially abundant microbes between the disease and control groups are included in the KG, based on values in the *“FDR”* column (representing false discovery rate) that are less than 0.05. These values indicate statistical significance, as determined using the Microbiome Multivariable Association with Linear Models (MaAsLin2) method^61^.

#### 4.2.10. Comparative Toxicogenomics Database (CTD)

The latest release (January 2025) of chemical-disease associations was downloaded from the CTD data downloads page (*“CTD_chemicals_diseases.tsv.gz”*). The *“DiseaseID”* column contains Medical Subject Headings (MeSH) IDs that are mapped to MONDO terms using the *“xref”* fields of the MONDO OWL file. Chemicals are extracted from the *“CasRN”* field or the *“ChemicalID”* field, which returns CAS-RN or MeSH IDs, respectively. Those IDs are mapped to ChEBI IDs using the Node Normalization API v1.4, which returns equivalent identifiers of chemicals from a variety of ontologies or standardized resources (Table 1). Chemicals are linked to diseases via the *“biolink:associated_with”* predicate.

### 4.3. Butyrate Analysis

We used a metagenomic analysis by Vital et al. of 3,184 sequenced bacterial genomes from the publicly available Integrated Microbial Genome database, which documented 225 predicted butyrate producing bacteria at the species or strain level. 216 of these taxa were able to be mapped to a NCBI Taxonomy ID using a pattern-based fuzzy string matching. First a direct string match was attempted. Next, brackets were included around the first term, assumed to be a genus name, and the match was attempted again. Finally *“sp.”* was included in the name and the match was attempted again. Any microbes that were not able to be mapped to an NCBI Taxonomy term were manually curated when possible, leaving only 3 *“not found”* taxa from the Vital et al. set.

We compared these taxa to those represented in KG-Microbe from various sources. Butyrate production is represented in the graph from BacDive or as functional annotations from UniProt. In BacDive, microbes are cited as being capable of producing butyrate from literature curation or in house experiments. One semantic representation involving UniProt data was microbes with proteins involved in biochemical reactions (Rhea nodes) that have butyrate as a product. Because reaction annotations in UniProt are undirected, we applied the eQuilibrator methodology to predict the direction of a given reaction, which uses group contribution to estimate free energy of formation (ΔfG°)^70^. Overall, this approach contributed 3,176 total taxa to the set of predicted butyrate producers from the KG. Another semantic representation involving UniProt data was microbes capable of defined sets of EC numbers, allowing for one missing annotated reaction. These EC sets were identical to those used in the Vital et al. metagenomic analysis to represent four butyrate producing pathways (the acetyl-CoA, glutarate, 4-aminobutyrate, and lysine pathways, Figure 3A).

We searched for all strains and species for relevant families using DuckDB over KG-Microbe-Biomedical-Function. We use the *“biolink:subclass_of”* edge to find all children of each given taxa. Because there exists taxa without a rank assignment in NCBI Taxonomy, we excluded these and any children of the taxa without rank from analysis. All species and strain level butyrate producers in KG-Microbe-Function were found, and their corresponding family and phylum assignment was identified in order to constrain the list by only those phyla relevant to the human gut^2^. We then evaluated key families known for butyrate production in the gut microbiome: Ruminococcaceae, Lachnospiraceae, Bacteroidaceae, Prevotellaceae, Clostridiaceae, Oscillospiraceae, and Eubacteriaceae^30^. Of that list, we evaluated the proportion of total taxa in the family, according to knowledge represented in the KG from NCBI Taxonomy, to butyrate producers (Figure 4C).

### 4.4. Disease Analysis

Microbes with a disease relationship were found in KG-Microbe-Biomedical-Function based on edge and node type. The edges “*biolink:associated_with_decreased_likelihood_of*” and “*biolink:associated_with_increased_likelihood_of*” were used for the analysis in both IBD and PD. To identify microbes involved in IBD, we included the diseases Crohn’s disease and Ulcerative colitis in the search, since they are children of the IBD disease class according to MONDO. The resulting microbe:disease pairs were sourced from Disbiome and Wallen et al. Microbes with an inconsistent direction of their disease relationship across studies were excluded. We then traversed KG-Microbe-Biomedical-Function, similarly to the methods described in the “Butyrate Annotation” section to find all strains or species of a query taxon, at any rank, at a finer taxonomic resolution. When no strains existed in NCBI Taxonomy beneath the given taxon, we evaluated species. From the collection of protein annotations to these taxa, we then characterized them as being butyrate producers or not based on the KG organismal and genomic trait representation in Figure 3A. We then compared the fraction of taxa annotated to produce butyrate out of all representative strains (or species) between those that increase or decrease likelihood of the disease.

### 4.5. Temperature Classification Model

A catboost multiclass classification model was trained using KG-Microbe EC and Rhea UniProt binary annotations. Rows and columns with less than two observations were removed. All four temperature classes were used, all with at least 40 training examples: psychrophilic, mesophilic, thermophilic, and hyperthermophilic. 25 EC features were removed because of identity with another EC feature. Data was split with stratification as 70/20/10% for train/validation/test, with hyperthermophiles having the fewest test data examples with 8. Training data consisted of 5,628 taxa and 6,177 EC and Rhea features. The catboost parameters were 200,000 iterations, depth of 4, learning rate of 0.05, L2 regularization term of 4, bagging temperature of 1, random strength of 6, a MultiClass loss function, random seed of 42, verbosity of 100, early stopping rounds of 50, and use best model set to True. CatBoost model SHAP values were computed using the TreeExplainer shap_values() method from the SHAP python package.

## 5. Data Availability

All KG resources are available in KG-Registry^12^. In addition, KG-Microbe-Core and KG-Microbe-Biomedical are available in KG-Hub. The code for the framework to create each transformed module for KG-Microbe graphs and to merge the non-function graphs can be accessed online at https://github.com/Knowledge-Graph-Hub/kg-microbe. The code to generate raw files for incorporating UniProt annotations can be accessed online at https://github.com/Knowledge-Graph-Hub/uniprot2s3. The code to merge transformed modules for the function graphs in the graph can be accessed online at https://github.com/Knowledge-Graph-Hub/kg-microbe-merge. The code for all analyses can be accessed online at https://github.com/bsantan/kg-microbe-paper.

## References

1. Kim, N. et al. Genome-resolved metagenomics: a game changer for microbiome medicine. Exp. Mol. Med. 56, 1501–1512 (2024).

2. Hou, K. et al. Microbiota in health and diseases. Signal Transduct. Target. Ther. 7, 135 (2022).

3. Langille, M. G. I. et al. Predictive functional profiling of microbial communities using 16S rRNA marker gene sequences. Nat. Biotechnol. 31, 814–821 (2013).

4. Mann, E. R., Lam, Y. K. & Uhlig, H. H. Short-chain fatty acids: linking diet, the microbiome and immunity. Nat. Rev. Immunol. 24, 577–595 (2024).

5. Putman, T. E. et al. The Monarch Initiative in 2024: an analytic platform integrating phenotypes, genes and diseases across species. Nucleic Acids Res. 52, D938–D949 (2024).

6. Callahan, T. J. et al. An open source knowledge graph ecosystem for the life sciences. Sci. Data 11, 363 (2024).

7. Ma, C., Liu, S. & Koslicki, D. MetagenomicKG: a knowledge graph for metagenomic applications. Preprint at 10.1101/2024.03.14.585056 (2024).

8. Goetz, S. L., Glen, A. K. & Glusman, G. MicrobiomeKG: Bridging Microbiome Research and Host Health Through Knowledge Graphs. Preprint at 10.1101/2024.10.10.617697 (2024).

9. Liu, T., Lan, G., Feenstra, K. A., Huang, Z. & Heringa, J. Towards a knowledge graph for pre-/probiotics and microbiota–gut–brain axis diseases. Sci. Rep. 12, 18977 (2022).

10. Liu, T. et al. Predicting the relationships between gut microbiota and mental disorders with knowledge graphs. Health Inf. Sci. Syst. 9, 3 (2021).

11. Unni, D. R. et al. Biolink Model: A universal schema for knowledge graphs in clinical, biomedical, and translational science. Clin. Transl. Sci. 15, 1848–1855 (2022).

12. KG-Registry

13. Schoch, C. L. et al. NCBI Taxonomy: a comprehensive update on curation, resources and tools. Database 2020, baaa062 (2020).

14. Söhngen, C., Bunk, B., Podstawka, A., Gleim, D. & Overmann, J. BacDive—the Bacterial Diversity Metadatabase. Nucleic Acids Res. 42, D592–D599 (2014).

15. Cébron, A. et al. BactoTraits – A functional trait database to evaluate how natural and man-induced changes influence the assembly of bacterial communities. Ecol. Indic. 130, 108047 (2021).

16. Madin, J. S. et al. A synthesis of bacterial and archaeal phenotypic trait data. Sci. Data 7, 170 (2020).

17. Reimer, L. C. et al. Bac *Dive* in 2022: the knowledge base for standardized bacterial and archaeal data. Nucleic Acids Res. 50, D741–D746 (2022).

18. Koblitz, J. et al. Media *Dive* : the expert-curated cultivation media database. Nucleic Acids Res. 51, D1531–D1538 (2023).

19. The UniProt Consortium et al. UniProt: the Universal Protein Knowledgebase in 2023. Nucleic Acids Res. 51, D523–D531 (2023).

20. Morgat, A. et al. UniPathway: a resource for the exploration and annotation of metabolic pathways. Nucleic Acids Res. 40, D761–D769 (2012).

21. Vasilevsky, N. A., et al. Mondo: Unifying Diseases for the World, by the World. http://medrxiv.org/lookup/doi/10.1101/2022.04.13.22273750 (2022) doi:10.1101/2022.04.13.22273750.

22. Janssens, Y. et al. Disbiome database: linking the microbiome to disease. BMC Microbiol. 18, 50 (2018).

23. Wallen, Z. D. et al. Metagenomics of Parkinson’s disease implicates the gut microbiome in multiple disease mechanisms. Nat. Commun. 13, 6958 (2022).

24. Davis, A. P. et al. Comparative Toxicogenomics Database (CTD): update 2021. Nucleic Acids Res. 49, D1138–D1143 (2021).

25. Peng, L., Li, Z.-R., Green, R. S., Holzmanr, I. R. & Lin, J. Butyrate Enhances the Intestinal Barrier by Facilitating Tight Junction Assembly via Activation of AMP-Activated Protein Kinase in Caco-2 Cell Monolayers. J. Nutr. 139, 1619–1625 (2009).

26. Recharla, N., Geesala, R. & Shi, X.-Z. Gut Microbial Metabolite Butyrate and Its Therapeutic Role in Inflammatory Bowel Disease: A Literature Review. Nutrients 15, 2275 (2023).

27. Beber, M. E. et al. eQuilibrator 3.0: a database solution for thermodynamic constant estimation. Nucleic Acids Res. 50, D603–D609 (2022).

28. Vital, M., Howe, A. C. & Tiedje, J. M. Revealing the Bacterial Butyrate Synthesis Pathways by Analyzing (Meta)genomic Data. mBio 5, e00889–14 (2014).

29. Markowitz, V. M. et al. IMG: the integrated microbial genomes database and comparative analysis system. Nucleic Acids Res. 40, D115–D122 (2012).

30. Singh, V. et al. Butyrate producers, “The Sentinel of Gut”: Their intestinal significance with and beyond butyrate, and prospective use as microbial therapeutics. Front. Microbiol. 13, 1103836 (2023).

31. Elford, J. D., Becht, N., Garssen, J., Kraneveld, A. D. & Perez-Pardo, P. Buty and the beast: the complex role of butyrate in Parkinson’s disease. Front. Pharmacol. 15, 1388401 (2024).

32. Rup, L. The Human Microbiome Project. Indian J. Microbiol. 52, 315–315 (2012).

33. Louca, S., Mazel, F., Doebeli, M. & Parfrey, L. W. A census-based estimate of Earth’s bacterial and archaeal diversity. PLOS Biol. 17, e3000106 (2019).

34. Lundberg, S. & Lee, S.-I. A Unified Approach to Interpreting Model Predictions. Preprint at 10.48550/ARXIV.1705.07874 (2017).

35. Chattopadhyay, M. K. et al. Increase in Oxidative Stress at Low Temperature in an Antarctic Bacterium. Curr. Microbiol. 62, 544–546 (2011).

36. Ladenstein, R. & Antranikian, G. Proteins from hyperthermophiles: Stability and enzymatic catalysis close to the boiling point of water. in Biotechnology of Extremophiles (ed. Antranikian, G.) vol. 61 37–85 (Springer Berlin Heidelberg, Berlin, Heidelberg, 1998).

37. Li, B., Kurihara, S., Kim, S. H., Liang, J. & Michael, A. J. A polyamine-independent role for *S*-adenosylmethionine decarboxylase. Biochem. J. 476, 2579–2594 (2019).

38. Becskei, A. & Rahaman, S. The life and death of RNA across temperatures. Comput. Struct. Biotechnol. J. 20, 4325–4336 (2022).

39. Parks, D. H. et al. GTDB: an ongoing census of bacterial and archaeal diversity through a phylogenetically consistent, rank normalized and complete genome-based taxonomy. Nucleic Acids Res. 50, D785–D794 (2022).

40. Kanehisa, M., Furumichi, M., Sato, Y., Kawashima, M. & Ishiguro-Watanabe, M. KEGG for taxonomy-based analysis of pathways and genomes. Nucleic Acids Res. 51, D587–D592 (2023).

41. Yao, G. et al. MicroPhenoDB Associates Metagenomic Data with Pathogenic Microbes, Microbial Core Genes, and Human Disease Phenotypes. Genomics Proteomics Bioinformatics 18, 760–772 (2020).

42. Olson, R. D. et al. Introducing the Bacterial and Viral Bioinformatics Resource Center (BV-BRC): a resource combining PATRIC, IRD and ViPR. Nucleic Acids Res. 51, D678–D689 (2023).

43. Santangelo, B., Bada, M., Hunter, L. & Lozupone, C. Hypothesizing Mechanistic Links between Microbes and Disease Using Knowledge Graphs. http://biorxiv.org/lookup/doi/10.1101/2023.12.01.569645 (2023) doi:10.1101/2023.12.01.569645.

44. Caspi, R. et al. The MetaCyc database of metabolic pathways and enzymes - a 2019 update. Nucleic Acids Res. 48, D445–D453 (2020).

45. Norsigian, C. J. et al. BiGG Models 2020: multi-strain genome-scale models and expansion across the phylogenetic tree. Nucleic Acids Res. gkz1054 (2019) doi:10.1093/nar/gkz1054.

46. Moretti, S., Tran, V. D. T., Mehl, F., Ibberson, M. & Pagni, M. MetaNetX/MNXref: unified namespace for metabolites and biochemical reactions in the context of metabolic models. Nucleic Acids Res. 49, D570–D574 (2021).

47. Heinken, A. et al. Genome-scale metabolic reconstruction of 7,302 human microorganisms for personalized medicine. Nat. Biotechnol. 1–12 (2023) doi:10.1038/s41587-022-01628-0.

48. Diener, C., Gibbons, S. M. & Resendis-Antonio, O. MICOM: Metagenome-Scale Modeling To Infer Metabolic Interactions in the Gut Microbiota. mSystems 5, e00606–19 (2020).

49. Santangelo, B. E. et al. Integrating biological knowledge for mechanistic inference in the host-associated microbiome. Front. Microbiol. 15, 1351678 (2024).

50. Caufield, J. H. et al. KG-Hub—building and exchanging biological knowledge graphs. Bioinformatics 39, btad418 (2023).

51. Weinstein, M. P. & Lewis, J. S. The Clinical and Laboratory Standards Institute Subcommittee on Antimicrobial Susceptibility Testing: Background, Organization, Functions, and Processes. J. Clin. Microbiol. 58, e01864–19 (2020).

52. bacteria-archaea-traits.

53. Degtyarenko, K. et al. ChEBI: a database and ontology for chemical entities of biological interest. Nucleic Acids Res. 36, D344–D350 (2007).

54. Buttigieg, P. et al. The environment ontology: contextualising biological and biomedical entities. J. Biomed. Semant. 4, 43 (2013).

55. Cooper, L. & Jaiswal, P. The Plant Ontology: A Tool for Plant Genomics. in Plant Bioinformatics (ed. Edwards, D.) vol. 1374 89–114 (Springer New York, New York, NY, 2016).

56. Mungall, C. J., Torniai, C., Gkoutos, G. V., Lewis, S. E. & Haendel, M. A. Uberon, an integrative multi-species anatomy ontology. Genome Biol. 13, R5 (2012).

57. Dooley, D. M. et al. FoodOn: a harmonized food ontology to increase global food traceability, quality control and data integration. Npj Sci. Food 2, 23 (2018).

58. PATO: Phenotypic Quality Ontology.

59. Jackson, R. C. et al. ROBOT: A Tool for Automating Ontology Workflows. BMC Bioinformatics 20, 407 (2019).

60. Ontology Access Kit (OAK).

61. Mallick, H. et al. Multivariable association discovery in population-scale meta-omics studies. PLOS Comput. Biol. 17, e1009442 (2021).

62. Hastings, J. et al. ChEBI in 2016: Improved services and an expanding collection of metabolites. Nucleic Acids Res. 44, D1214–D1219 (2016).

63. Jacobs, A. et al. CAS Common Chemistry in 2021: Expanding Access to Trusted Chemical Information for the Scientific Community. J. Chem. Inf. Model. 62, 2737–2743 (2022).

64. Kim, S. et al. PubChem 2023 update. Nucleic Acids Res. 51, D1373–D1380 (2023).

65. Bairoch, A. The ENZYME database in 2000. Nucleic Acids Res. 28, 304–305 (2000).

66 . The Gene Ontology Consortium. The Gene Ontology Resource: 20 years and still GOing strong. Nucleic Acids Res. 47, D330–D338 (2019).

67. Köhler, S. et al. The Human Phenotype Ontology in 2021. Nucleic Acids Res. 49, D1207–D1217 (2021).

68. Seal, R. L. et al. Genenames.org: the HGNC resources in 2023. Nucleic Acids Res. 51, D1003–D1009 (2023).

69. Bansal, P. et al. Rhea, the reaction knowledgebase in 2022. Nucleic Acids Res. 50, D693–D700 (2022).

70. Haghbakhsh, R., Raeissi, S. & Duarte, A. R. C. Group contribution and atomic contribution models for the prediction of various physical properties of deep eutectic solvents. Sci. Rep. 11, 6684 (2021).

