## Supplemental Figure 1 for "KG-Microbe - Building Modular and Scalable Knowledge Graphs for Microbiome and Microbial Sciences"

**Supplementary Figure 1**

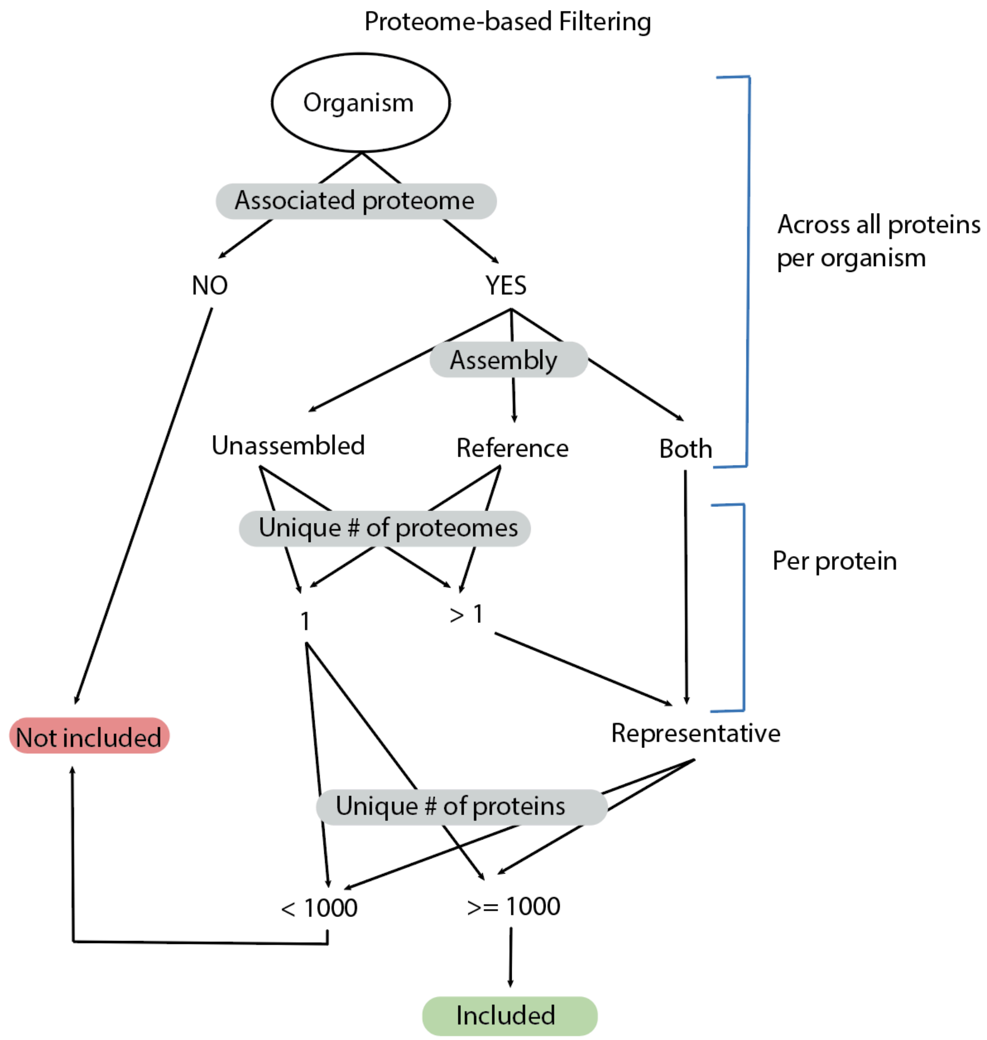


The KG-Microbe strain-level pan-proteome approach, which uses all representative proteins available for a given NCBI Taxonomy species. Each organism is represented as an NCBI Taxonomy identifier in the UniProt database, and those are found via the uniprot2s3 repository. Only those with associated proteomes are included in the transform. Proteins are marked as Unassembled or part of the Reference proteome, in which case Reference assemblies are prioritized if present. Only one is taken when more than one entry exists per protein to avoid duplication. Finally, only proteomes for organisms that have more than 1000 proteins are included.
